# Erythroblast-derived mediators program neutrophil development and function

**DOI:** 10.64898/2026.03.15.711869

**Authors:** Duco S. Koenis, Roberta de Matteis, Esteban A. Gomez, Antal Rot, Jesmond Dalli

**Affiliations:** William Harvey Research Institute, Barts and The London, Faculty of Medicine and Dentistry, Queen Mary University of London, Charterhouse Square, London, EC1M 6BQ UK; Centre for Inflammation and Therapeutic Innovation, Queen Mary University of London, London, UK

**Author notes:** Department of Biostatistics & Health Informatics, Institute of Psychiatry Psychology & Neuroscience, King’s College London, 16 De Crespigny Park, London SE5 8AB, UK. Corresponding author: Prof Jesmond Dalli Ph.D, William Harvey Research Institute, John Vane Science Centre, Charterhouse Square, London. EC1M 6BQ.

**Keywords:** neutrophil development, erythroblast-derived mediators, specialized pro-resolving mediators, Alox15 enzyme, granulopoiesis

## Abstract

Granulopoiesis is a tightly regulated process encompassing the production, maturation, and release of neutrophils in the bone marrow, ensuring their optimal physiological contribution to host defence. The mechanisms that maintain a balanced regulation of neutrophil effector functions during this process remain incompletely understood.

Here, we identify bone marrow-resident erythroblasts as a key source of specialized pro-resolving mediators (SPMs) and show that they imprint neutrophil development and function. Terminally differentiating erythroblasts highly express the key SPM biosynthetic enzyme 12/15-lipoxygenase (Alox15) and accordingly generate several SPMs, including n-3 docosapentaenoic acid-derived Resolvin D5 (RvD5_n-3 DPA_). Conditional erythroblast-specific depletion of Alox15 decreased bone marrow SPM levels and caused altered neutrophil phenotypes including augmented release of reactive oxygen species and neutrophil extracellular traps and increased migration to chemotactic stimuli, leading to increased neutrophil sequestration in peripheral organs and concomitant neutropenia. Mice with erythroblast-specific Alox15 depletion displayed impaired bacterial clearance in experimental peritonitis and increased severity in DSS-colitis. Aberrant neutrophil phenotypes were rectified by the reconstitution of RvD5_n-3 DPA_ and were largely recapitulated by the specific depletion of the SPM receptor Gpr101 in bone marrow macrophages, but not in neutrophil precursors, suggesting the involvement of the macrophage niche in mediating the SPM effects on granulopoiesis.

Our findings establish a central role for erythroblasts and their SPM production in instructing granulopoiesis for balanced functional neutrophil responses.

**Key Points:** Erythroblasts are a source of Alox15-dependent SPMs that regulate BM erythroblastic island integrity and neutrophil maturation and function

Loss of erythroblast Alox15 disrupts neutrophil development and function, defects that are restored add-back of RvD5_n-3 DPA_.

## Introduction

Neutrophils are essential for host defence and tissue homeostasis. Traditionally viewed as the body’s first line of defence against bacterial and fungal pathogens^1^, they are now recognized as multifunctional cells that also contribute to tissue repair and regeneration, promoting collagen deposition, angiogenesis, and restoration of barrier integrity^2,3^. In humans and some other mammals, neutrophils are the most abundant immune cell type, reflecting their fundamental role in maintaining physiological balance. Because of their short lifespan, neutrophils must be continually replenished through granulopoiesis, a tightly regulated process that preserves both neutrophil cell numbers and functions^1,4^.

Mature circulating neutrophils are terminally differentiated cells that have limited capacity to change their transcriptome or profoundly alter the bias of functional responses after exiting the bone marrow. Therefore, neutrophil imprinting that equips them to meet the organism’s peripheral needs occurs within the bone marrow. During neutrophil development their phenotypes and functions are shaped by a combination of soluble growth factors and cytokines (i.e. G-CSF, GM-CSF, IL-6, CXCL12, TNF, type I/II IFNs), metabolic and soluble mediators (i.e. retinoic acid and ROS), and cellular niche partners (including mesenchymal stromal cells, osteoblasts, endothelial cells, macrophages, megakaryocytes and sympathetic nerves)^5–14^. Together, these signals reprogram intrinsic transcriptional and epigenetic pathways, imprinting neutrophil functional states that prime the cells for potent antimicrobial defence, activities that are effective against invading pathogens but can, in some circumstances, also cause collateral tissue damage. Conversely, remarkably little is known about the bone marrow mechanisms that regulate neutrophils and counterbalance their aggressive phenotypes to achieve homeostatic functional balance under steady-state conditions. Disbalanced neutrophil development can lead to their numerical changes and qualitative defects, contributing to infection susceptibility, chronic inflammation, and impaired tissue repair^2^.

Among the pathways that remain poorly understood in this context are those governed by specialized pro-resolving mediators (SPMs), a family of bioactive lipid mediators enzymatically derived from omega-3 fatty acids. SPMs are best known for orchestrating the resolution of inflammation, promoting tissue repair, and restraining emergency myelopoiesis^15–19^. Notably, dysregulated SPM biosynthesis has been documented in multiple chronic disease settings, including ageing, where bone-marrow myeloid output is perturbed and pathogenic phagocyte phenotypes emerge. Although SPMs are present in the bone marrow under steady-state conditions^20^, their potential role in programming neutrophil development during homeostatic granulopoiesis remains entirely unexplored.

Using a systematic approach coupling transcriptomic approaches with lipid mediator profiling and novel transgenic models, herein we identify erythroblast progenitors as a previously unrecognized prime source of SPMs that program neutrophil development in the bone marrow. We show that erythroblast-derived SPMs, synthesized via 12/15-lipoxygenase (*Alox15*), are essential for maintaining marrow SPM tone and instructing balanced neutrophil maturation. Loss of erythroblast Alox15 results in disrupted granulopoiesis and aberrant neutrophil activation, while restitution with the Alox15 product Resolvin(Rv)D5_n-3 DPA_ re-balances neutrophil homeostasis. These findings reveal a previously unappreciated erythroblast– neutrophil axis that links lipid mediator signalling to hematopoietic programming, uncovering a new layer through which resolution pathways sustain immune equilibrium.

## Methods

This section briefly describes the methods; further details are provided in the supplemental Methods.

### In vivo experiments

All in vivo experiments were conducted in Specific Opportunistic Pathogen Free facilities at Charles River Laboratories (UK). Mice were maintained on a 12 h light/dark cycle with ad libitum access to food and water, and were randomly assigned to control or treatment groups. All procedures were approved by the UK Home Office (PP5449341) and performed in accordance with the Animals (Scientific Procedures) Act (1986) and LASA welfare guidelines.

### Statistical analysis

Data are presented as mean ± SEM and analysed using GraphPad Prism v10.

Statistical tests, group sizes, and replicate numbers are detailed in figure legends. Normality was assessed by Shapiro–Wilk testing. Parametric or non-parametric tests were applied as appropriate, with p < 0.05 considered significant.

## Results

### Alox15 expression in erythroblasts is critical for maintaining bone marrow SPM levels

In peripheral tissues, Alox15 is a key enzyme in the biosynthesis of specialised pro-resolving mediators (SPMs) that drive the resolution of inflammation. We therefore first asked whether Alox15 plays a comparable role in regulating SPM production within the bone marrow. Lipid mediator profiling revealed that, consistent with previous reports, SPMs are abundant in the bone marrow of wild-type mice^20^. Notably, global deletion of Alox15 resulted in a profound reduction or complete loss of multiple SPM species, establishing Alox15 as a central determinant of bone-marrow SPM availability (Figure.1A,B and Figure.S1).

**Figure 1:**
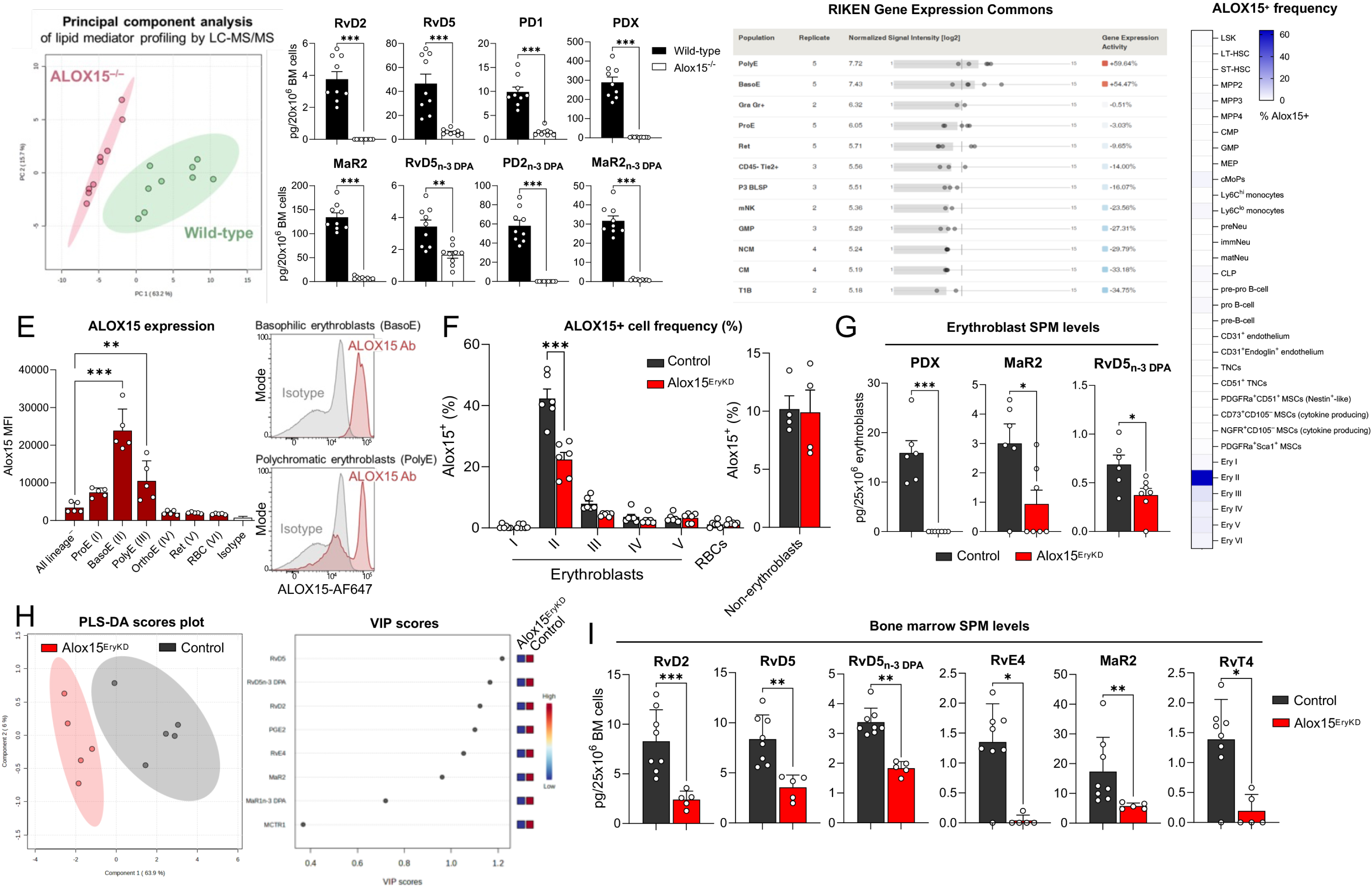
Alox15 expression in erythroblasts is critical for maintaining bone marrow SPM levels. (A,B) Lipid mediator profiling of bone marrow from Alox15-deficient mice (Alox15^-/-^) and Alox15-sufficient wild-type mice; (a) PCA score plot; (b) levels of individual SPM, RvD = D-series Resolvins, PD = Protectins, MaR = Maresins, RvD_n-3 DPA_ = n-3 DPA-derived D-series Resolvins, MaR_n-3 DPA_ = n-3 DPA-derived Maresins, PD_n-3 DPA_ = n-3 DPA-derived Protectins. n= 9 mice per group. (C) *Alox15* transcript expression in various bone marrow hematopoietic cell types from the RIKEN Gene Expression Commons database. (D) Frequency of ALOX15^+^ cells in various bone marrow cell populations from wild-type mice as assessed by intracellular flow cytometry. n = 6 mice. (E) Median fluorescence intensity (MFI) of ALOX15 staining in erythroblast populations from wild-type mice as assessed by intracellular flow cytometry. n = 5 mice. (F) Frequency of ALOX15^+^ erythroblast populations from erythroblast-specific Alox15-depleted mice (Hbb:cre-Alox15^fl/fl^; abbreviated as Alox15^EryKD^), compared to Alox15-sufficient littermates (Alox15^fl/fl^; abbreviated as Control). Inset shows frequency of ALOX15^+^ non-erythroblast cell lineages. n = 6 mice per group. (G) SPM levels in Ter119^+^ erythroblasts enriched from Alox15^EryKD^ mice and Alox15-sufficient Alox15^fl/fl^ (Control) littermate bone marrow. n = 7 Alox15^EryKD^ mice, 6 Control littermates. (H-I) Lipid mediator profiling of bone marrow from Alox15^EryKD^ mice compared to Alox15-sufficient littermates (Control); (h) Partial Least Squares-Discriminant Analysis (PLS-DA) and Variable Importance in Projection (VIP) score analysis of LM profiling dataset; (i) Quantification of select SPMs with VIP score > 1.0; n = 8 Control, 5 Alox15^EryKD^ mice. Data are shown as mean ± SEM. Statistical significance was determined using Mann-Whitney U-test (b,i), one-way ANOVA (e) or two-way ANOVA (g) with Holm-Sidak multiple comparison correction. *p <0.05, **p <0.01, ***p <0.001 as indicated.

We next sought to identify the cellular sources of Alox15 responsible for SPM biosynthesis in this compartment. Analysis of the RIKEN Gene Expression Commons, a public database of experimentally validated gene expression levels across a wide range of tissues^21^, revealed striking enrichment of Alox15 transcripts in basophilic erythroblasts (BasoE) and polychromatic erythroblasts (PolyE) compared to all other mouse bone marrow cell types (Figure.1C). hese findings were validated experimentally by intracellular flow cytometry, which demonstrated high levels of Alox15 protein expression selectively within BasoE and PolyE subsets, with minimal expression detected in other haematopoietic lineages (Figure.1D,E, Figure.S2-5).

Given this prominent erythroblast-restricted expression, we investigated whether erythroblasts contribute directly to bone-marrow SPM production. To address this, we generated mice with erythroblast-specific depletion of Alox15 by crossing *loxP*-flanked *Alox15* expressing mice (Alox15^fl/fl^)^22^ with heterozygote mice expressing Cre recombinase under the control of the human beta hemoglobin (HBB) promoter (Hbb:cre)^23^. Assessment of Alox15 expression in erythroblasts from these transgenic erythroblast-specific Alox15 depleted (Alox15^EryKD^) mice by ICFC showed a roughly 50% reduction in erythroblasts staining positive for Alox15 compared to Alox15-sufficient (Alox15^fl/fl^) control littermates (Figure.1F).

We next examined whether these cells produced SPMs. Using LC–MS/MS we profiled lipid mediators in erythroid cells isolated from the bone marrow of Alox15^EryKD^ mice and Alox15-sufficient control littermates. We identified multiple Alox15-derived SPMs that included the docosahexaenoic acid-derived Protectin D1 (PD1) and Maresin (MaR)2, as well as the RvD5_n-3 DPA_, which were significantly reduced in erythroblasts from Alox15^EryKD^ mice (Figure.1G, Figure.S2). Furthermore, profiling of the entire bone marrow of these Alox15^EryKD^ mice showed a marked shift in SPM levels (Figure.1H), largely driven by significant reductions in the levels of the SPMs RvD2, RvD5, RvD5_n-3 DPA_, RvE4, MaR2, and RvT4 (Figure.1I).

Taken together, these results show that Alox15 expression in erythroblasts is important for the production of SPMs within the bone marrow niche.

### Erythroblast-derived Alox15 governs neutrophil maturation and effector programming

Given that SPM levels were markedly reduced in Alox15^EryKD^ mice, we next evaluated the functional impact that loss of this mediator has on haematopoiesis. To this end, we performed single-cell RNA sequencing (scRNA-seq) on whole bone marrow from Alox15^EryKD^ mice and Alox15-sufficient control littermates, generating a dataset of 26,468 cells encompassing all major haematopoietic populations. Comparative analysis revealed shifts in population frequencies between genotypes, indicating altered bone-marrow organization (Figure.2A,B).

**Figure 2:**
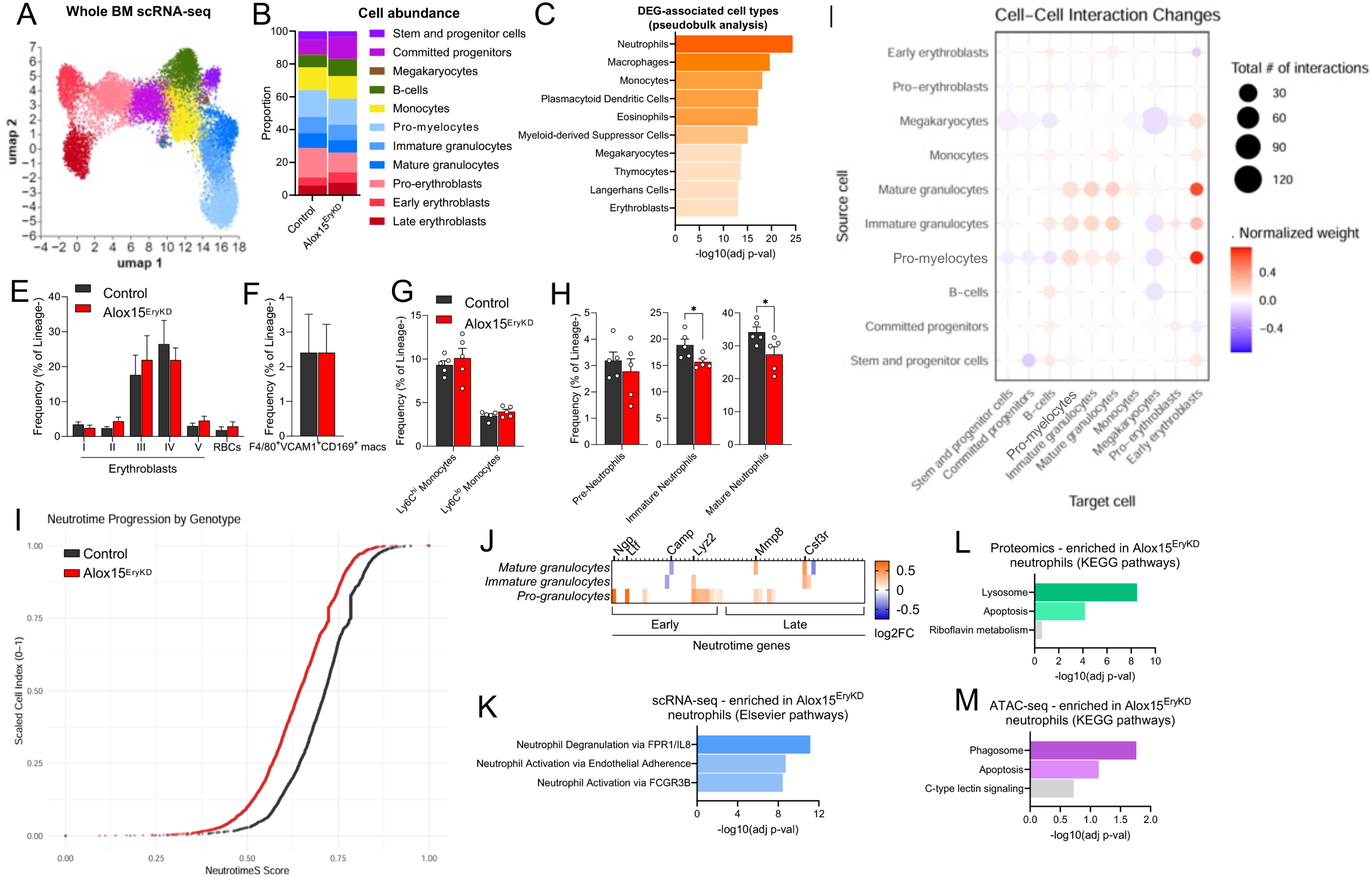
Single-cell RNA sequencing analysis reveals defective neutrophil maturation and effector function transcriptional programs in mice with erythroblast-specific depletion of Alox15. (aA-D) Single-cell RNA sequencing (scRNA-seq) on whole bone marrow from Alox15^EryKD^ mice and Alox15-sufficient Alox15^fl/fl^ littermates (control); (A) UMAP projection of 26,468 cells total, of which 13,973 Alox15^EryKD^ cells and 12,495 Control cells; (B) Proportion of each identified cell population based on marker gene expression and their frequency in the scRNA-seq dataset; (C) Cell types associated with differentially-expressed genes from pseudobulk analysis between all Alox15^EryKD^ and control cells using PanglaoDB annotation; (D) Cell-cell interaction analysis using LIgand-receptor ANalysis framework (LIANA) showing interactions increased (orange) or decreased (blue) in Alox15^EryKD^ bone marrow compared to control bone marrow based on ligand-receptor pair transcripts in the scRNA-seq dataset. n = 6 mice per group. (E-H) Frequency of various hematopoietic cell subsets in bone marrow from Alox15^EryKD^ mice and Alox15-sufficient Alox15^fl/fl^ littermates (control); (E) nucleated erythroblast subsets and red blood cells (RBC); (F) marrow niche macrophages; (G) monocytes; (H) neutrophils. n = 5 mice per group. (I-J) Neutrophil maturation (Neutrotime) analysis of granulocyte subpopulations in the scRNA-seq dataset; (I) Neutrotime-S score for Alox15^EryKD^ and control granulocyte subpopulations; (J) log2 fold-change in significantly differentially-expressed (adjusted p-value < 0.05) Neutrotime signature genes between Alox15^EryKD^ and Control pro-, immature, and mature granulocyte populations. Positive values indicates higher expression in Alox15^EryKD^ mice, negative values indicate higher expression in Alox15-sufficient Alox15^fl/fl^ littermate Controls. (K) EnrichR Elsevier pathway analysis of differentially-expressed genes between Alox15^EryKD^ and Control granulocyte populations. (L) KEGG pathway analysis of genes associated with significantly increased accessible chromatin sites as determined by ATAC-seq in Alox15^EryKD^ bone marrow neutrophils compared to Control neutrophils. n = 4 mice per group. (m) KEGG pathway analysis of significantly increased protein peptides in Alox15^EryKD^ bone marrow neutrophils compared to Control neutrophils. n = 4 mice per group. Data are shown as mean ± SEM. Statistical significance was determined using Mann-Whitney U-test (F,H), or two-way ANOVA (E,G) with Holm-Sidak multiple comparison correction. *p <0.05, **p <0.01, ***p <0.001 as indicated.

To identify the cell types most affected by erythroblast Alox15 depletion, we conducted pseudo-bulk differential gene expression analysis followed by annotation using PanglaoDB^24^ and Enrichr^25^ (Figure.2C). Unexpectedly, erythroblasts themselves were minimally affected, whereas the three most perturbed populations were exclusively myeloid (i.e. neutrophils, macrophages, and monocytes) suggesting indirect regulation of myelopoiesis by erythroblast-derived signals (Figure.2C). Consistent with this, cell–cell interaction analysis using LIgand-receptor ANalysis framework (LIANA)^26^, revealed pronounced alterations in predicted communication between early erythroblasts, which highly express Alox15 (Figure.1C), and pro-myelocytes and mature granulocytes (Figure.2D).

Guided by these findings, we quantified myeloid populations by flow cytometry. Frequencies of erythroblasts, bone-marrow niche macrophages, and Ly6C^hi^ and Ly6C^lo^ monocytes were unchanged (Figure.2E-G, Figure.S3,5). In contrast, both immature and mature neutrophil populations were significantly reduced in Alox15^EryKD^ mice, indicating a selective defect in granulopoiesis (Figure.2H). We therefore interrogated neutrophil transcriptional states in greater depth.

Using the Neutrotime^27^ maturation framework, we found a skewing of neutrophils towards a less mature transcriptional phenotype in Alox15^EryKD^ bone marrow (Figure.2I). This characterised by elevated expression of early maturation markers, including *Ngp*, *Ltf,* and *Lyz2*, alongside increased expression of *Csf3r*, which encodes for the critical neutrophil developmental regulator granulocyte colony-stimulating factor (G-CSF) receptor (Figure.2J). Pathway analysis on all granulocyte cells within the scRNA-seq dataset using Enrichr showed enrichment of genes associated with neutrophil degranulation, activation via endothelial adherence, and activation via FCGR3B (Figure.2K).

To determine whether these transcriptional changes were reflected at the protein level, we performed unbiased proteomic profiling of neutrophils. This revealed significant enrichment of proteins involved in phagolysosome formation and apoptotic execution (Figure.2L), indicating altered effector programming. To identify upstream regulatory mechanisms, we complemented these analyses with ATAC-seq. Neutrophils from Alox15^EryKD^ mice exhibited increased chromatin accessibility at loci associated with lysosomal pathways and apoptosis (Figure.2M), providing evidence for epigenetic priming of this altered state.

### Erythroblast Alox15 depletion induces hyperactive and functionally defective neutrophils

The in-depth molecular profiling of the chromatin accessibility, transcriptome and proteome of neutrophils from Alox15^EryKD^ bone marrow pointed towards an aberrant neutrophil phenotype. Therefore, we next evaluated whether these changes translated to altered cell functions in these cells. Neutrophils form a first line of defense to infection and injury through a combination of phagocytosis, degranulation, and the production of cytotoxic effectors such as neutrophil extracellular traps (NETs) and reactive oxygen species (ROS). We found that neutrophils isolated from Alox15^EryKD^ bone marrow produced significantly more ROS (Figure.3A) and NETs (Figure.3B) in response to sterile pro-inflammatory stimuli *in vitro*, while being significantly less effective in phagocytosis of gram-negative *E. coli* and fungal zymosan particles (Figure.3C) compared to neutrophils from Alox15-sufficient control littermates. Notably, phagocytosis of gram-positive *S. aureus* particles was unchanged from control littermates (Figure.3D). In addition to these effector functions, we found that neutrophils isolated from Alox15^EryKD^ bone marrow displayed enhanced chemotaxis in response to formyl-methionine tripeptide (fMLF) *in vitro* (Figure.3D).

**Figure 3:**
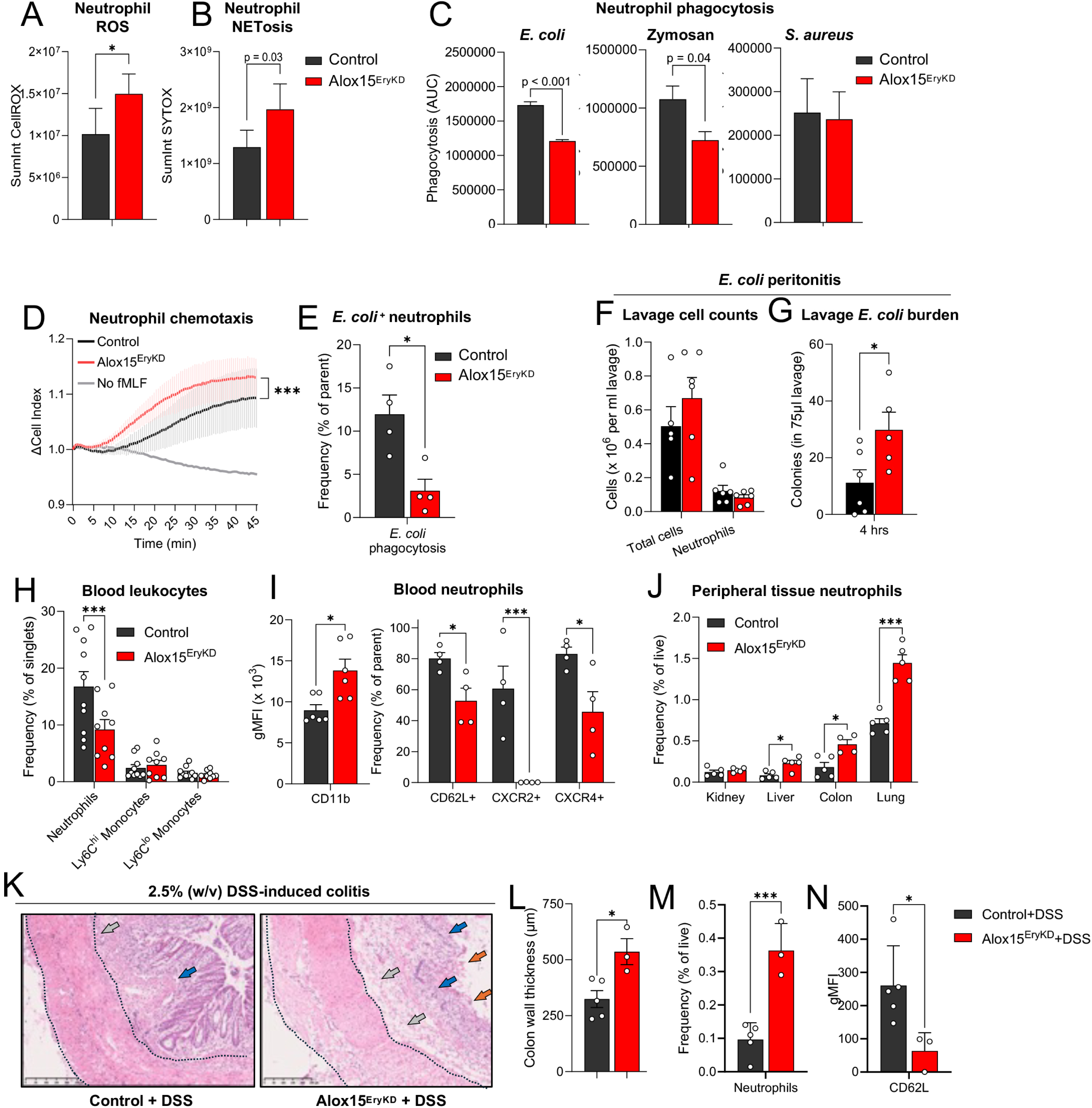
Altered neutrophil development and function in mice with erythroblast-specific depletion of Alox15. (A-D) Functional assays on neutrophils enriched from Alox15^EryKD^ mice and Alox15-sufficient Alox15^fl/fl^ (Control) littermate bone marrow; (A) reactive oxygen species (ROS) production in response to N-formyl-methionyl-leucyl-phenylalanine (fMLF); (B) neutrophil extracellular trap (NET) production in response to phorbol 12-myristate 13-acetate (PMA); (C) phagocytosis of pHrodo-labelled *E. coli*, Zymosan, or *S. aureus* bioparticles; (D) chemotaxis towards fMLF. N = 5 Control littermates, 4 Alox15^EryKD^ mice. (E) Uptake of pHrodo-labelled *E. coli* bioparticles by circulating neutrophils from Alox15^EryKD^ mice and Alox15-sufficient Alox15^fl/fl^ (Control) littermates. n = 4 mice per group. (F-G) *In vivo* mouse *E. coli* peritonitis model in which mice were injected with 10^5^ CFU *E. coli*; (F) peritoneal lavage total cell and neutrophil counts at 4 hours post-infection; (G) *E. coli* colony outgrowth at 4 hours post-infection. n = 6 mice per group. (H-I) Circulating leukocyte phenotyping in whole blood from Alox15^EryKD^ mice and Alox15-sufficient Alox15^fl/fl^ (Control) littermates at baseline; (H) Leukocyte frequencies, n = 9 mice per group; (I) Neutrophil cell surface marker expression, n = 4 mice per group. (J) Neutrophil abundance in digested peripheral tissues from Alox15^EryKD^ mice and Alox15-sufficient Alox15^fl/fl^ (Control) littermates . n = 5 mice per group. (K-N) 2.5% (w/v) Dextran sulphate sodium (DSS)-induced colitis in Alox15^EryKD^ mice and Alox15-sufficient Alox15^fl/fl^ (Control) littermates; (K) Representative images of H&E stained distal colon sections, arrows indicate loss of crypt architecture (blue), epithelial erosion (orange), and increased mucosal remodeling (gray), dashed line demarcates colon wall thickness, scale bar = 250 µm; (L) quantification of colon wall thickness from H&E stained sections described in K; (M) frequency of neutrophils in digested distal colon tissues; (N) geometric mean fluorescence intensity (gMFI) of CD62L staining on neutrophils from digested distal colon tissues. n = 5 Control DSS-treated mice, n = 3 Alox15^EryKD^ DSS-treated mice. Data are shown as mean ± SEM. Statistical significance was determined using Mann-Whitney U-test (a-c,e,g,m,n), one-way ANOVA (N-P) or two-way ANOVA (F, H, J) with Holm-Sidak multiple comparison correction. *p <0.05, **p <0.01, ***p <0.001 as indicated.

Given the dysregulated phenotype of bone marrow neutrophils in Alox15^EryKD^ mice, we next examined whether these abnormalities persisted after their release into circulation. Circulating neutrophils from Alox15^EryKD^ mice exhibited significantly impaired phagocytosis of Gram-negative *E. coli* particles in *ex vivo* whole-blood assays (Figure.3E, Figure.S7). Moreover, *in vivo* challenge with *E. coli* revealed a marked defect in bacterial clearance at 4 hours post-infection, despite comparable neutrophil recruitment to the peritoneum in Alox15^EryKD^ mice and Alox15-sufficient control littermates (Figure.3F,G). These findings indicate that functional defects in neutrophils of Alox15^EryKD^ mice persist beyond the bone marrow, potentially compromising their capacity for effective host defense.

Together with these functional defects, Alox15^EryKD^ mice displayed a substantial reduction in circulating neutrophil numbers, with neutrophil frequency in blood decreased by approximately 50% relative to Alox15-sufficient littermate controls (Figure.3H). This reduction in circulating neutrophils was more pronounced than the 20% reduction in mature neutrophil frequency observed in the bone marrow of these mice (Figure.2H), with many of the Alox15^EryKD^ mice showing neutropenia (defined as <500 neutrophils per microliter blood^28^) at baseline (Figure.S8A). Furthermore, upon investigation of surface markers on circulating leukocytes using flow cytometry we found that neutrophils from Alox15^EryKD^ mice showed significantly higher surface expression of the β₂-integrin CD11b and concomitant downregulation of L-selectin (CD62L) and the chemokine receptors CXCR2 and CXCR4 (Figure.3I). This pattern is consistent with an activated neutrophil phenotype in which selectin shedding and chemokine-receptor internalization accompany integrin upregulation^29^.

Given the activated surface marker phenotype on circulating neutrophils, we speculated that the exacerbated decline in circulating versus bone marrow neutrophils in Alox15^EryKD^ mice could be due to abnormal sequestration of these cells in peripheral tissue vascular beds. Indeed, we found increased neutrophil numbers in the lung, liver, and colon of Alox15^EryKD^ mice, while neutrophil numbers in the kidney were comparable to Alox15-sufficient control littermates (Figure.3J, Figure.S9). Taken together, these observations indicate that aberrant programming of Alox15^EryKD^ neutrophils in the bone marrow drives their premature activation and pathological retention within mucosal and hepatic tissues.

To further assess the pathological consequences of the heightened activation state and increased neutrophil presence in the colon of Alox15^EryKD^ mice, we induced sterile colitis by administering dextran sodium sulfate (DSS) in the drinking water. Distal colons from Alox15^EryKD^ mice exhibited exacerbated inflammation, characterized by loss of crypt architecture, epithelial erosion, and mucosal remodeling (Figure.3L). Quantification of the colon wall thickness showed a significant increase in DSS-treated Alox15^EryKD^ mice compared to DSS-treated Alox15-sufficient control littermates (Figure.3M), while total colon length was not significantly different (Figure.S8B). On the other hand, neutrophils were significantly more abundant in the distal colon of Alox15^EryKD^ mice (Figure.3N), and these cells expressed significantly lower levels of CD62L compared to Alox15-sufficient control littermates (Figure.3O).

Together, these results indicate that depletion of Alox15 in erythroblasts compromises neutrophil programming, predisposing them to excessive activation and inflammatory tissue damage.

### RvD5_n-3_ DPA reverses neutrophil defects in Alox15^EryKD^ mice

Given that erythroblast-specific Alox15 depletion reduced bone marrow SPM levels and disrupted neutrophil development and function, we next sought to determine whether restoring specific SPMs could reverse these abnormalities. We focused on RvD5_n-3 DPA_, an Alox15-derived mediator whose levels were markedly decreased in erythroblasts from Alox15^EryKD^ mice as well as bone marrow from these mice (Figure.1G,I). We therefore performed RvD5_n-3 DPA_ add-back experiments to test whether replenishing this mediator could rescue the neutrophil defects observed in Alox15^EryKD^ mice.

Administration of RvD5_n-3 DPA_ over a two-week period to Alox15^EryKD^ mice via intravenous injection (Figure.4A) led to a significant increase in mature neutrophil frequency in the bone marrow (Figure.4B and Figure.S3) and circulation (Figure.4C) when compared to vehicle treated Alox15^EryKD^ mice. Moreover, neutrophils isolated from the bone marrow of Alox15^EryKD^ mice treated with RvD5_n-3 DPA_ were better at phagocytosing *E. coli* particles (Figure.4D) and zymosan particles (Figure.4E), while production of NETs in response to phorbol 12-myristate 13-acetate (PMA) was normalized back to levels observed in Alox15-sufficient control littermates (Figure.4F). On the other hand, *in vivo* RvD5_n-3 DPA_ dosing had no effect on the heightened ROS production observed in Alox15^EryKD^ mice (Figure.4G), suggesting that this biological response was under the control of other Alox15-derived SPM or mechanisms.

**Figure 4:**
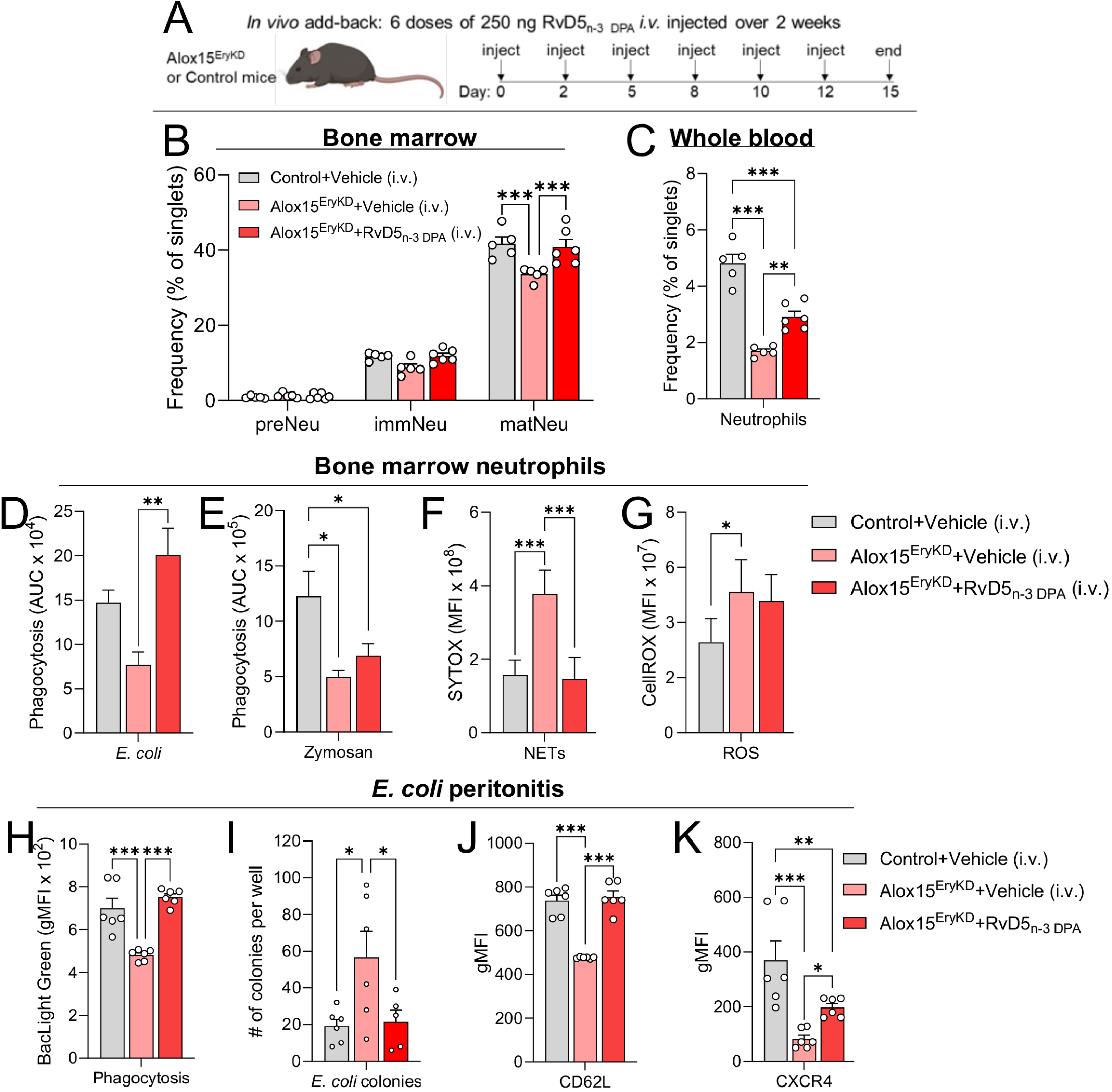
RvD5_n-3 DPA_ add-back partially corrects neutrophil abnormalities in erythroblast-specific Alox15-depletion mice at baseline and during infection. *In vivo* add-back of the SPM RvD5_n-3 DPA_ in Alox15^EryKD^ mice and Alox15-sufficient Alox15^fl/fl^ (Control) littermates via intravenous (i.v.) injection; (A) schematic overview of add-back regimen; (B) frequency of pre-neutrophils (preNeu), immature neutrophils (immNeu), and mature neutrophils (matNeu) in the bone marrow of treated mice at day 15 post-treatment initiation; (C) frequency of neutrophils in the circulation of treated mice at day 15; (d-g) phagocytosis of pHrodo-labeled *E. coli* (D) or Zymosan bioparticles (e) *in vitro* by neutrophils isolated from the bone marrow of *in vivo* treated mice at day 15 post-treatment initiation; (F) production of neutrophil extracellular traps (NETs) in response to PMA and (G) production of reactive oxygen species (ROS) in response to fMLF by neutrophils isolated from the bone marrow of *in vivo* treated mice at day 15. n = 5 Control vehicle-treated mice, n = 5 Alox15^EryKD^ vehicle-treated mice, n = 6 Alox15^EryKD^ RvD5_n-3 DPA_-treated mice. (H-K) *In vivo* add-back of the SPM RvD5_n-3 DPA_ in Alox15^EryKD^ and Control via intravenous (i.v.) injection followed by inoculation of 10^5^ CFU live fluorescently labelled *E. coli*; After 4h peritoneal lavages were collected and (H) peritoneal lavage neutrophil phagocytosis of live *E. coli* was determined using flow cytometry; (I) *E. coli* colony counts in cultured peritoneal lavage at 4 hrs post-infection; (J-K) cell surface expression of (J) CD62L and (K) CXCR4 on peritoneal lavage neutrophils was determined using flow cytometry. n = 6 mice per group. Data are shown as mean ± SEM. Statistical significance was determined using one-way ANOVA (D-M) or two-way ANOVA (C) with Holm-Sidak multiple comparison correction. *p <0.05, **p <0.01, ***p <0.001 as indicated.

**Figure 5:**
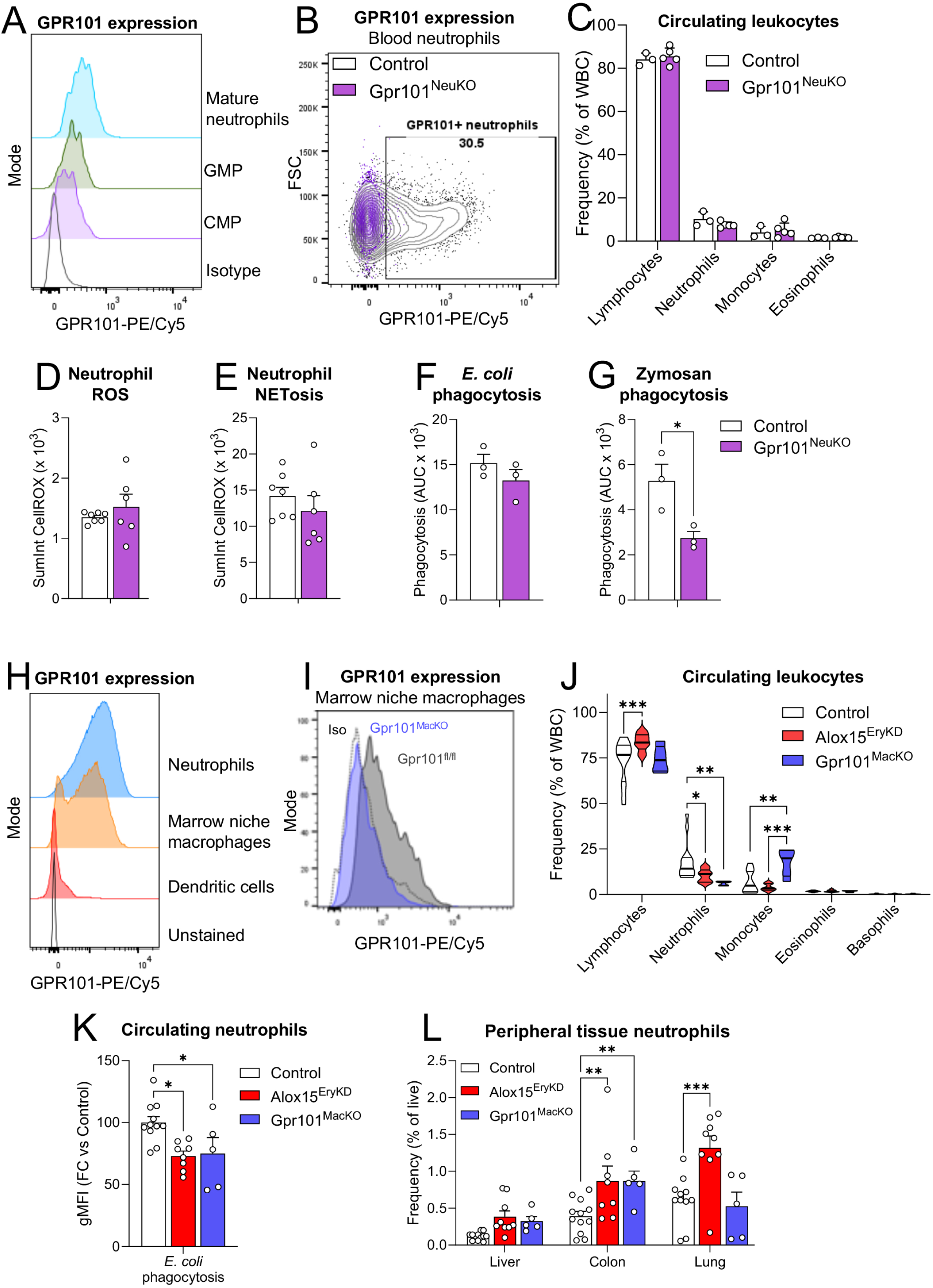
Deletion of the RvD5_n-3 DPA_ receptor GPR101 in mononuclear phagocytes, but not in neutrophils, partially recapitulates aberrant neutrophil phenotypes observed in erythroblast-specific Alox15-depletion mice. (A-B) Representative histograms of Gpr101 expression: (A) on mature neutrophils and GMP and CMP neutrophil progenitors from wild-type mouse bone marrow; (B) on circulating neutrophils from granulocyte-specific Gpr101 deficient Ms4a3:cre-Gpr101^fl/fl^ mice (Gpr101^NeuKO^) or Gpr101-sufficient Gpr101^fl/fl^ (Control) littermates. (C) Circulating leukocyte frequencies in Gpr101^NeuKO^ mice or control (Gpr101^fl/fl^) littermates from Procyte complete blood count analyzer. n = 3 Control, 5 Gpr101^NeuKO^ mice. (D-G) Neutrophil *in vitro* functional assays using purified bone marrow neutrophils from Gpr101^NeuKO^ mice or control (Gpr101^fl/fl^) littermates; (D) ROS production; (E) NET production; (F) pHrodo-labeled *E. coli* phagocytosis; (G) pHrodo-labeled zymosan bioparticle phagocytosis. n = 6 mice per group for D-E; n = 3 mice per group for F-G. (H) Representative histograms of Gpr101 expression on wild-type bone marrow Ly6G^+^CD11b^+^ mature neutrophils, F4/80^+^VCAM1^+^CD169^+^ marrow niche macrophages, or Gr-1^–^B220^–^CD1c^+^ dendritic cells. (I) Representative histograms of Gpr101 expression on marrow niche macrophages from wild-type control or mononuclear phagocyte-specific Gpr101 deficient Cx3cr1:cre-Gpr101^fl/fl^ mice (Gpr101^MacKO^). (J) Circulating leukocyte frequencies in whole blood from Alox15^EryKD^, Gpr101^MacKO^, or littermate Control mice. n = 15 Control, 9 Alox15^EryKD^, 5 Gpr101^MacKO^ mice. (K) Uptake of pHrodo-labelled *E. coli* bioparticles by circulating neutrophils in ex-vivo incubations of whole blood from Alox15^EryKD^, Gpr101^MacKO^, or littermate Control mice. n = 11 Control, 8 Alox15^EryKD^, 5 Gpr101^MacKO^ mice. (L) Neutrophil abundance in digested peripheral tissues from Alox15^EryKD^, Gpr101^MacKO^, or littermate Control mice. n = 12 Control, 9 Alox15^EryKD^, 5 Gpr101^MacKO^ mice. Data are shown as histograms normalized to mode (A, H, J), as mean ± SEM (C-G, K, I), or as median with quartiles (J). Statistical significance was determined using Mann-Whitney U-test (D-G), or one-way ANOVA (K) or two-way ANOVA (C, J, L) with Holm-Sidak multiple comparison correction. *p <0.05, **p <0.01, ***p <0.001 as indicated.

We next evaluated whether *in vivo* administration of RvD5_n-3 DPA_ normalized neutrophil function during infection. In a model of peritoneal infection with live *E. coli*, intravenous RvD5_n-3 DPA_ treatment enhanced neutrophil phagocytosis of live bacteria (Figure.4H), resulting in improved bacterial containment and clearance, as evidenced by a significant reduction in peritoneal exudate bacterial burden (Figure.4I). Moreover, RvD5_n-3 DPA_ dosing restored the surface expression of CD62L and CXCR4 on circulating neutrophils from Alox15^EryKD^ mice to levels comparable to those of Alox15-sufficient control littermates (Figure.4J,K), indicating a reprogramming of the neutrophil activation state toward homeostasis.

Together, these findings demonstrate that *in vivo* administration of RvD5_n-3 DPA_ restores both functional competence and phenotypic balance of neutrophils in Alox15^EryKD^ mice.

### GPR101 expression in progenitors and macrophages drives RvD5_n-3 DPA_–dependent neutrophil programming

Since *in vivo* administration of RvD5_n-3 DPA_ restored neutrophil phenotype and function in Alox15^EryKD^ mice, we next asked whether this mediator directly regulates neutrophil programming by acting on neutrophil progenitors. To address this, we first examined expression of the RvD5_n-3 DPA_ receptor, GPR101, which is known to be present on mature neutrophils^30^. We found that GPR101 is also expressed across all neutrophil progenitor subsets (Figure.5A, Figure.S10), suggesting a potential role for this receptor during neutrophil maturation.

To test the functional relevance of GPR101 in neutrophil programming under homeostatic conditions, we generated a granulocyte-specific Gpr101 knockout mouse by crossing *Ms4a3*:cre mice (in which Cre recombinase is expressed under the granulocyte/monocyte-restricted *Ms4a3* gene promoter^31^) with Gpr101^fl/fl^ mice. The resulting *Ms4a3*:cre-Gpr101^fl/fl^ mice, termed Gpr101^NeuKO^, lacked GPR101 expression in neutrophils (Figure.5B). Unlike Alox15^EryKD^ mice, these animals displayed normal circulating neutrophil numbers (Figure.5C) and their bone marrow neutrophils did not exhibit elevated ROS or NET formation following inflammatory stimulation (Figure.5D,E). Interestingly, however, these neutrophils showed a selective defect in phagocytosis of zymosan, mirroring a phenotype observed in Alox15^EryKD^ mice, while phagocytosis of *E. coli* particles was unaffected (Figure.5F,G). Together, these findings suggest that, while GPR101 expression on neutrophil progenitors contributes to their steady-state programming, it does not fully account for the range of defects observed in Alox15^EryKD^ mice. Since much of the neutrophil phenotype Alox15^EryKD^ was restored by RvD5_n-3 DPA_ treatment, we hypothesized that additional cell types are involved in mediating the broader biological activities of this mediator.

Given that single-cell RNA-seq analysis of Alox15^EryKD^ bone marrow cells revealed marked alterations in macrophage transcriptional states (Figure.2C), combined with the fact that these cells closely interact with both erythroblasts and granulocytes within erythroblastic islands^32^ and can regulate granulopoiesis^13^, we speculated that bone marrow niche–resident macrophages might mediate the effects of RvD5_n-3 DPA_ on neutrophil programming. To test this hypothesis we explored GPR101 expression on bone marrow macrophages using flow cytometry, observing that these cells express this receptor (Figure.5H). We then generated a transgenic mouse line in which GPR101 is deleted in mononuclear phagocytes by crossing mice expressing Cre-recombinase under the control of the *Cx3cr1* gene promoter^33^ with *loxP-* flanked *Gpr101* expressing mice (Cx3cr1:cre-Gpr101^fl/fl^; abbreviated to Gpr101^MacKO^), which lacked expression of GPR101 on bone marrow niche-resident macrophages (Figure.5I). Since both *Ms4a3* and *Cx3cr1* can be expressed in certain dendritic cell subsets, we verified that GPR101 was not physiologically expressed on bone marrow dendritic cells (Figure.5H).

Strikingly, Gpr101^MacKO^ mice exhibited a marked reduction in circulating neutrophil frequencies, closely resembling the phenotype observed in Alox15^EryKD^ mice (Figure.5J). Circulating neutrophils from Gpr101^MacKO^ mice also displayed similarly impaired phagocytosis of *E. coli* particles as those from Alox15^EryKD^ mice when compared to cells from Gpr101 and Alox15-sufficient control littermates (Figure.5K). Moreover, we detected significant neutrophil accumulation in the colon and trends towards increases in the liver, but not in the lung, of Gpr101^MacKO^ mice, partially mirroring the tissue distribution seen in Alox15^EryKD^ mice (Figure.5L).

Together, these findings identify macrophage GPR101 expression as a key mediator of erythroblast-derived RvD5_n-3 DPA_ biology, linking marrow niche macrophage activity to the programming of neutrophil development and function.

### Expression of GPR101 on mononuclear phagocytes is required for rescue of neutrophil phenotypes by RvD5^n-3 DPA^

Given the emerging role of bone marrow niche macrophages in neutrophil programming, we next focused on a specialized niche where developing neutrophils, SPM-producing erythroblasts, and GPR101-expressing macrophages coexist in close proximity: the erythroblastic island (EBI)^32^. Using established imaging flow cytometry protocols^34^, we detected EBIs in the bone marrow of Gpr101^MacKO^, Alox15^EryKD^, and Alox15 and Gpr101-sufficient control littermate mice (Figure.6A and Figure.S11). In Alox15^EryKD^ bone marrow, these EBIs had significantly higher staining for Ly6G and S100A9 (Figure.6B), while Gpr101^MacKO^ EBIs only had higher staining for S100A9 (Figure.6C) compared to control littermates, suggestive of enhanced neutrophil presence in these EBIs. Furthermore, administration of RvD5_n-3 DPA_ over a two-week period to Alox15^EryKD^ mice via intravenous injection (as described in Figure.4) led to a normalization of Ly6G staining in Alox15^EryKD^ EBIs (Figure.6D). Notably, both Alox15^EryKD^ mice and Gpr101^MacKO^ bone marrow exhibited comparable reductions in cytokines critical for neutrophil development, namely G-CSF^1,12^ and IL-18^35,36^(Figure.6E). Taken together, these results suggest that a disruption in canonical EBI signaling centered on Gpr101-expressing macrophages contributes to defective neutrophil programming in Alox15^EryKD^ and Gpr101^MacKO^ mice.

**Figure 6:**
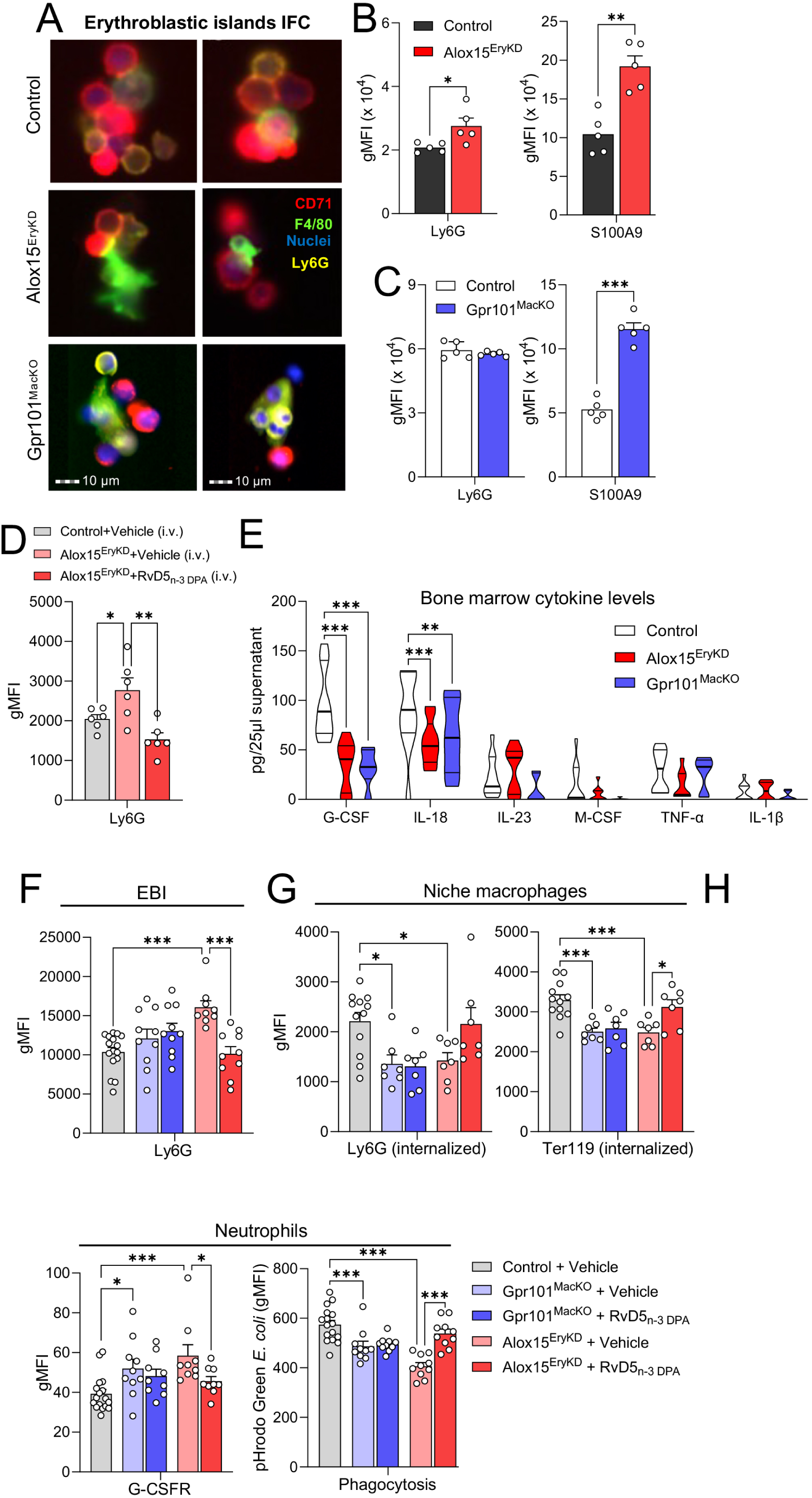
Expression of the RvD5_n-3 DPA_ receptor GPR101 on mononuclear phagocytes is required for rescue of neutrophil phenotypes by RvD5_n-3 DPA_. (A-C) ImageStream imaging flow cytometry (IFC) analysis of erythroblastic islands (EBI) in bone marrow from Alox15^EryKD^ mice, Gpr101^MacKO^ mice, or Alox15 and Gpr101-sufficient littermates (Control); (A) representative images, scale bar = 10 µm; (B) geometric mean fluorescence intensity (gMFI) of Ly6G and S100A9 neutrophil marker stainings in bone marrow EBIs from Control or Alox15^EryKD^ mice; (C) gMFI of Ly6G and S100A9 neutrophil marker stainings in bone marrow EBIs from Control or Gpr101^MacKO^ mice. n = 5 Alox15^fl/fl^ (Control), 5 Alox15^EryKD^ mice (for b); n = 5 Gpr101^fl/fl^ (Control), 5 Gpr101^MacKO^ mice (for c). (D) gMFI of Ly6G neutrophil marker staining in bone marrow EBIs at day 15 post-treatment initiation of *in vivo* RvD5_n-3 DPA_ add-back via intravenous (i.v.) injection in Alox15^EryKD^ and Control littermate mice as described in Fig. 4. n = 6 mice per group. (E) Bone marrow cytokine levels in Alox15^EryKD^ mice, Gpr101^MacKO^ mice, or Alox15 and Gpr101-sufficient littermates (Control). n = 11 Control, 9 Alox15^EryKD^, 6 Gpr101^MacKO^ mice. (F-H) *Ex vivo* cultures of intact femurs from Alox15^EryKD^ mice, Gpr101^MacKO^ mice, or Alox15 and Gpr101-sufficient littermates (Control) with or without 10 nM RvD5_n-3 DPA_ for 96 hours; (F) Ly6G expression in cultured femur bone marrow EBIs, n = 16 Control, 10 Gpr101^MacKO^, 10 Alox15^EryKD^ mice per group; (G) Uptake of Ter119^+^ and Ly6G^+^ particles by cultured femur bone marrow F4/80^+^VCAM1^+^CD169^+^ niche macrophages, n = 12 Control, 7 Gpr101^MacKO^, 7 Alox15^EryKD^ mice per group; (H) G-CSFR expression and phagocytosis of pHrodo-labeled *E. coli* by cultured femur bone marrow neutrophils, n = 16 Control, 10 Gpr101^MacKO^, 10 Alox15^EryKD^ mice per group. Data are shown as mean ± SEM (B-E, G-L) or median with quartiles (f). Statistical significance was determined using Mann-Whitney U-test (B-E), or two-way ANOVA (F-L) with Holm-Sidak multiple comparison correction. *p <0.05, **p <0.01, ***p <0.001 as indicated.

We next set out to establish both the requirement for RvD5_n-3 DPA_-Gpr101 signaling on niche macrophages as well as the possible role of EBI dysregulation in the defective neutrophil programming observed in Gpr101^MacKO^ and Alox15^EryKD^ mice. To that end, we cultured surgically opened, intact femurs from Gpr101^MacKO^ and Alox15^EryKD^ mice *ex vivo* in the presence of RvD5_n-3 DPA_ and compared them to femurs treated with vehicle or femurs from Gpr101 and Alox15-sufficient control littermates. We found that EBIs from Alox15^EryKD^ femurs had significantly higher Ly6G staining, consistent with *in vivo* observations (Figure.6B,D), which could be normalized to control littermate levels by *ex vivo* RvD5_n-3 DPA_ treatment (Figure.6F).

Since clearance of neutrophils by niche macrophages can trigger G-CSF production and is essential for maintaining neutrophil production^37^, we next assessed intracellular staining of neutrophil and erythroblast markers in niche macrophages as a readout for efferocytosis of apoptotic neutrophils and erythrophagocytosis, respectively. Consistent with previous observations that SPMs are crucial regulators of macrophage phagocytosis and efferocytosis^38^, we found that Gpr101^MacKO^ and Alox15^EryKD^ niche macrophages had reduced staining for ingested neutrophils and erythroid cells (Figure.6G). Notably, *ex vivo* treatment with RvD5_n-3 DPA_ could restore efferocytosis by Alox15^EryKD^ niche macrophages but not Gpr101^MacKO^ niche macrophages (Figure.6G).

Finally, RvD5_n-3 DPA_ treatment normalized G-CSF receptor (G-CSFR) expression and *E.* coli particle phagocytosis in neutrophils from *ex vivo* cultured Alox15^EryKD^ femurs, but not in neutrophils in cultures from Gpr101^MacKO^ mice (Figure.6H).

Taken together, these data establish a crucial role for GPR101-expressing macrophages in mediating the beneficial neutrophil reprogramming effects of RvD5_n-3 DPA_.

## Discussion

The present study identifies erythroblasts as a previously unrecognised source of specialised pro-resolving mediators (SPMs) that govern neutrophil programming within the bone marrow. We demonstrate that erythroblast expression of the SPM-biosynthetic enzyme Alox15 sustains local SPM production, preserves erythroblastic island (EBI) integrity, and supports balanced neutrophil maturation and effector function. Erythroblast-specific depletion of Alox15 markedly reduced marrow levels of RvD5_n-3 DPA_ and related SPMs, leading to defective neutrophil differentiation, impaired bacterial clearance, and aberrant sequestration of neutrophils in peripheral tissues. Restoration of RvD5_n-3 DPA_ normalised neutrophil output and function primarily through GPR101-dependent macrophage signalling, defining a lipid mediator axis that couples erythroid metabolism to granulopoiesis.

These findings redefine erythroblasts as immunoregulatory cells that coordinate erythroid and myeloid homeostasis. Beyond their established role in erythropoiesis, erythroblasts emerge here as metabolically active hubs that generate pro-resolving cues shaping developing immune lineages. Within EBIs, erythroblast-derived RvD5_n-3 DPA_ acts predominantly via GPR101-expressing niche macrophages to modulate neutrophil maturation. This cross-lineage communication introduces a new paradigm in which pro-resolving lipid mediator signalling is embedded within the bone marrow to calibrate neutrophil activation thresholds prior to their release into the circulation^14,39^. Such regulation complements cytokine- and transcription factor–driven granulopoietic programs, ensuring that emerging neutrophils are competent for host defence while restrained to limit collateral tissue injury. In this context, our findings extend emerging evidence that erythroid progenitors regulate immunity through cytokine production, chemokine retention, retention of chemokines by atypical chemokine receptor 1^40^ and expression of immunomodulatory receptors and ligands ^41,42^.

Mechanistically, Alox15 depletion in erythroblasts produced a profound imbalance in neutrophil programming. Single-cell transcriptomic and proteomic analyses revealed persistence of early differentiation signatures with enrichment of activation, degranulation, and oxidative pathways. Functionally, neutrophils displayed excessive reactive oxygen species and neutrophil extracellular trap formation, coupled with impaired phagocytosis and delayed apoptosis, features characteristic of inflammatory neutrophil subsets observed in chronic disease^43–45^. These changes were accompanied by reduced levels of G-CSF, IL-18, and M-CSF, cytokines critical for neutrophil, macrophage, and erythroid development, as well as disruption of canonical EBIs^13,35,36^. Together, these findings indicate that erythroblast-derived SPMs maintain both the structural and cytokine integrity of the granulopoietic niche. The resulting mismatch between activation and maturation provides a mechanistic explanation for the neutropenia and excessive tissue sequestration observed following erythroblast-specific Alox15 loss.

Genetic dissection of RvD5_n-3 DPA_ receptor pathways further defined the cellular hierarchy of this axis. Macrophage-specific deletion of Gpr101 recapitulated the major phenotypes of erythroblast Alox15 deficiency, including reduced circulating neutrophils, impaired phagocytosis, and increased tissue accumulation, establishing niche macrophages as a principal conduit of erythroblast-derived SPM signalling. Deletion of Gpr101 within the neutrophil lineage also produced measurable defects, indicating additional direct effects on maturing neutrophils. Thus, RvD5_n-3 DPA_ engages GPR101 both indirectly, through macrophage-mediated niche regulation, and directly within the neutrophil lineage to fine-tune activation and effector readiness. This stepwise mode of action extends the concept of resolution mediator biology to steady-state haematopoiesis and positions the RvD5_n-3 DPA_– GPR101 axis as a developmental checkpoint ensuring balanced neutrophil function before deployment.

Together, our results uncover a hierarchical lipid-mediator network in which erythroblast-derived SPMs coordinate the maturation and resolution programming of neutrophils. They also define a role for macrophage and neutrophil/neutrophil progenitor GPR101 in mediating, at least in part, this response by conveying the biological activities of RvD5_n-3 DPA_. This erythroblast–macrophage–neutrophil axis integrates metabolic, structural, and immunological control within the bone marrow, ensuring balanced neutrophil production and function. The identification of this pathway provides a conceptual framework for understanding how tissue-protective signalling is embedded in hematopoietic niches and suggests new therapeutic avenues: enhancing erythroblast SPM synthesis or selectively activating GPR101 could restore neutrophil homeostasis and resolution capacity in inflammatory and myelodysplastic diseases.

## Acknowledgements

This research was supported by funding from Barts Charity (grant no: MGU0439). This research was also supported by the CRUK FLOW CYTOMETRY CORE SERVICE GRANT at Barts Cancer Institute (CTRQQR-2021\100004) and made use of Queen Mary’s Apocrita HPC facility, supported by QMUL Research-IT (http://doi.org/10.5281/zenodo.438045).

## Authorship Contributions

D.K., A.R., and J.D. conceived and designed the experimental approaches described in the manuscript. J.D. conceived and directed the overall research plan. D.K., R.d.M., and E.A.G. performed the experiments. D.K. and J.D. drafted the manuscript, and all authors contributed to data interpretation, manuscript revision, and approved the final version.

---

**Figure S1:**
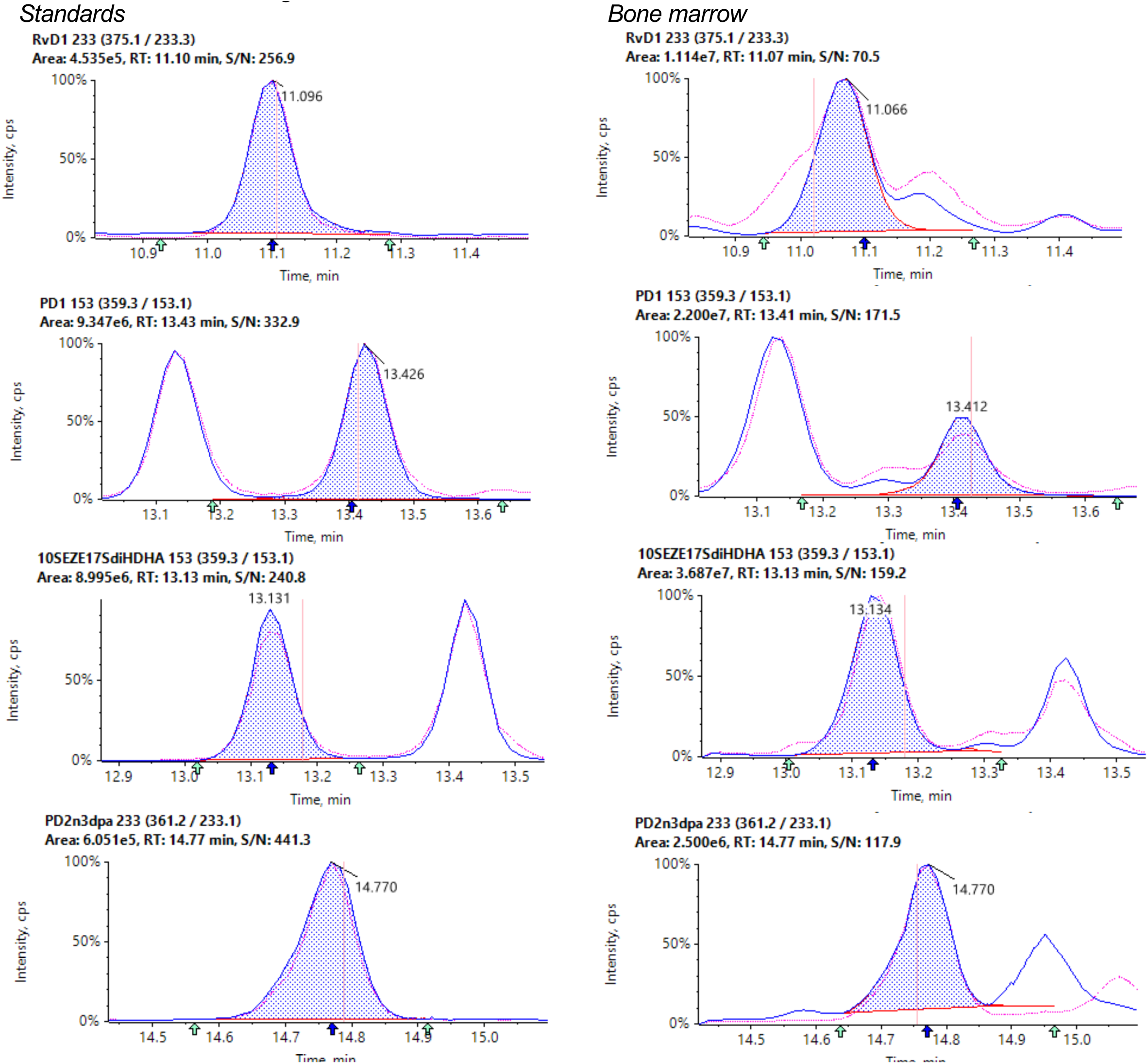
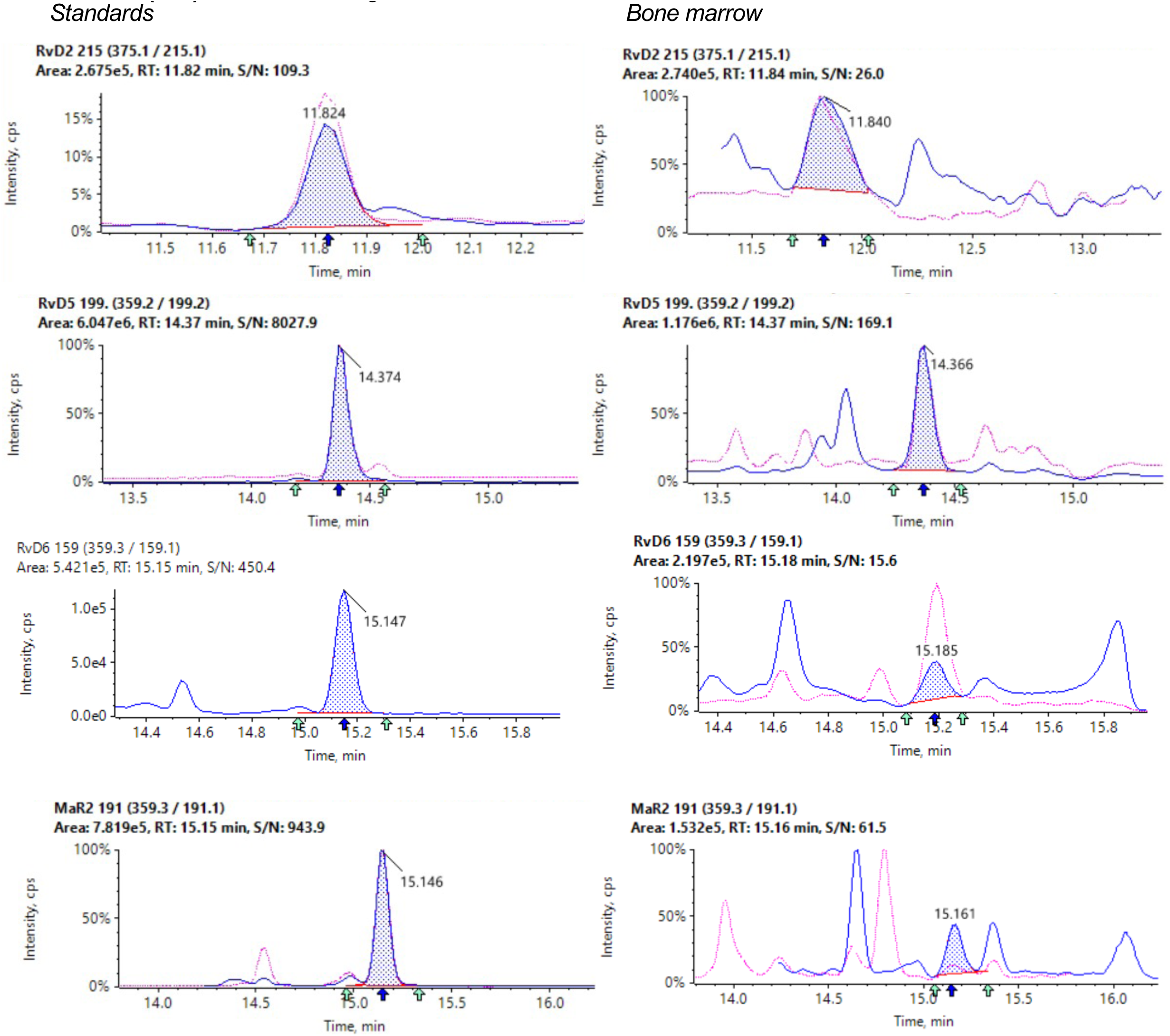
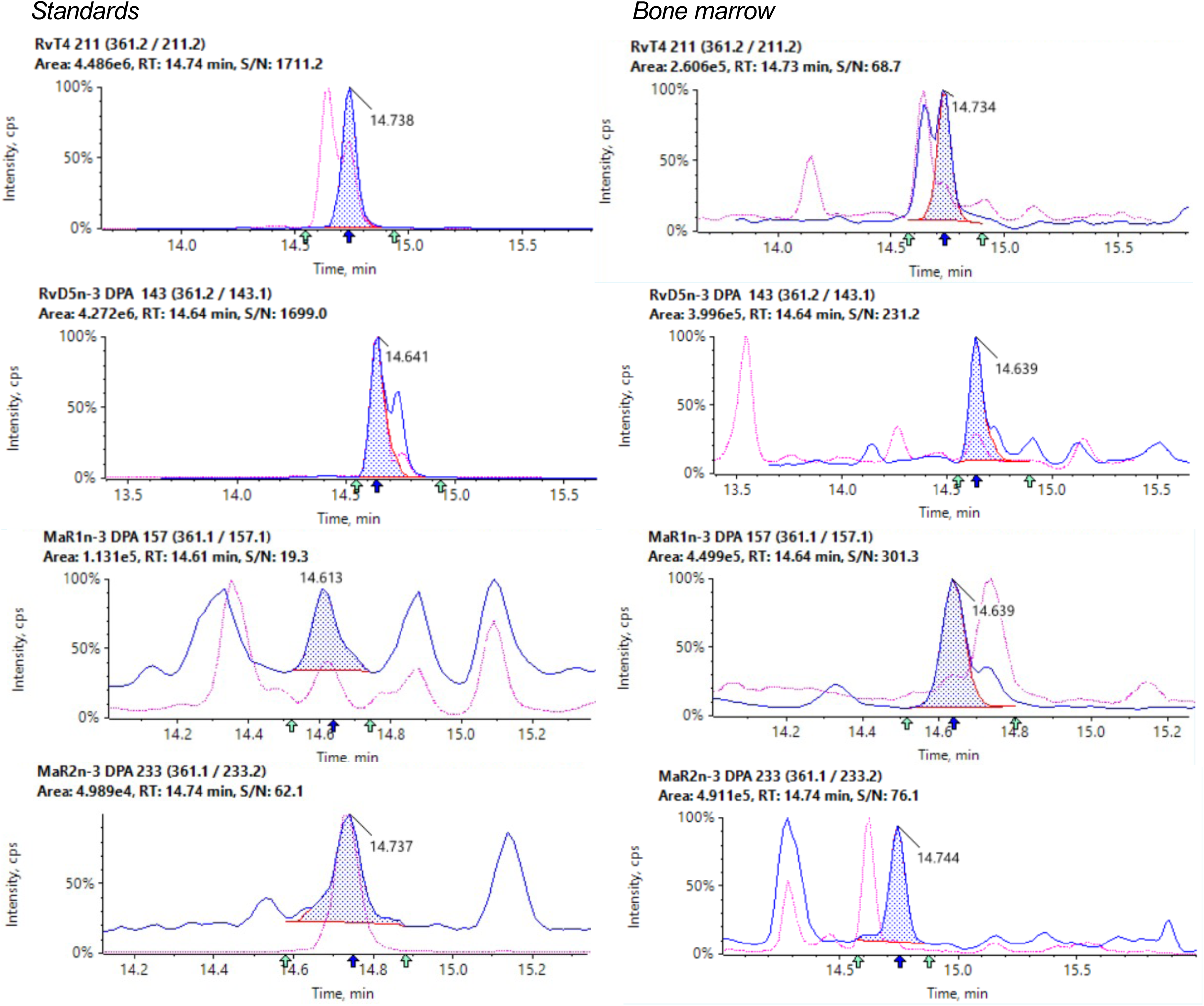
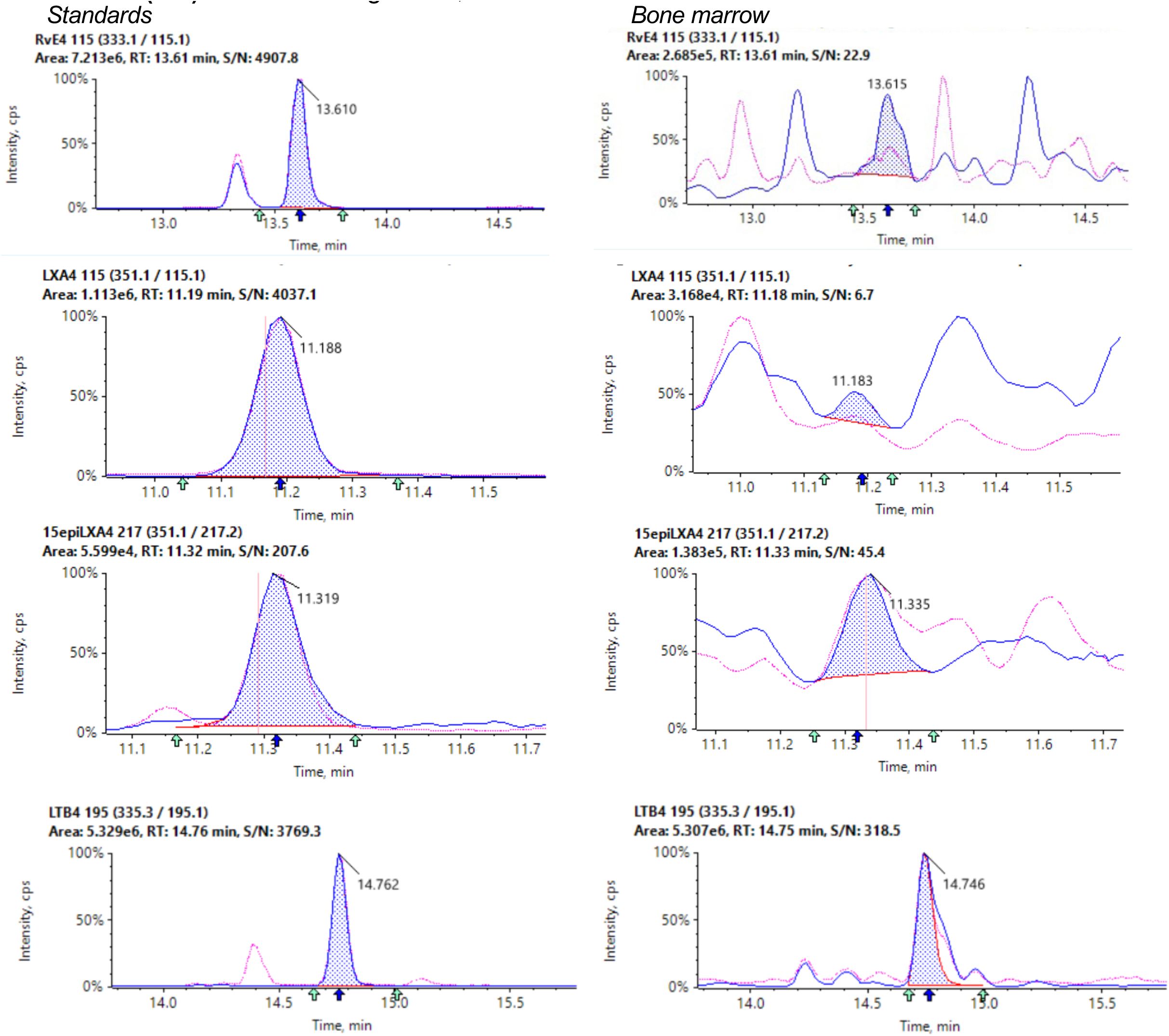
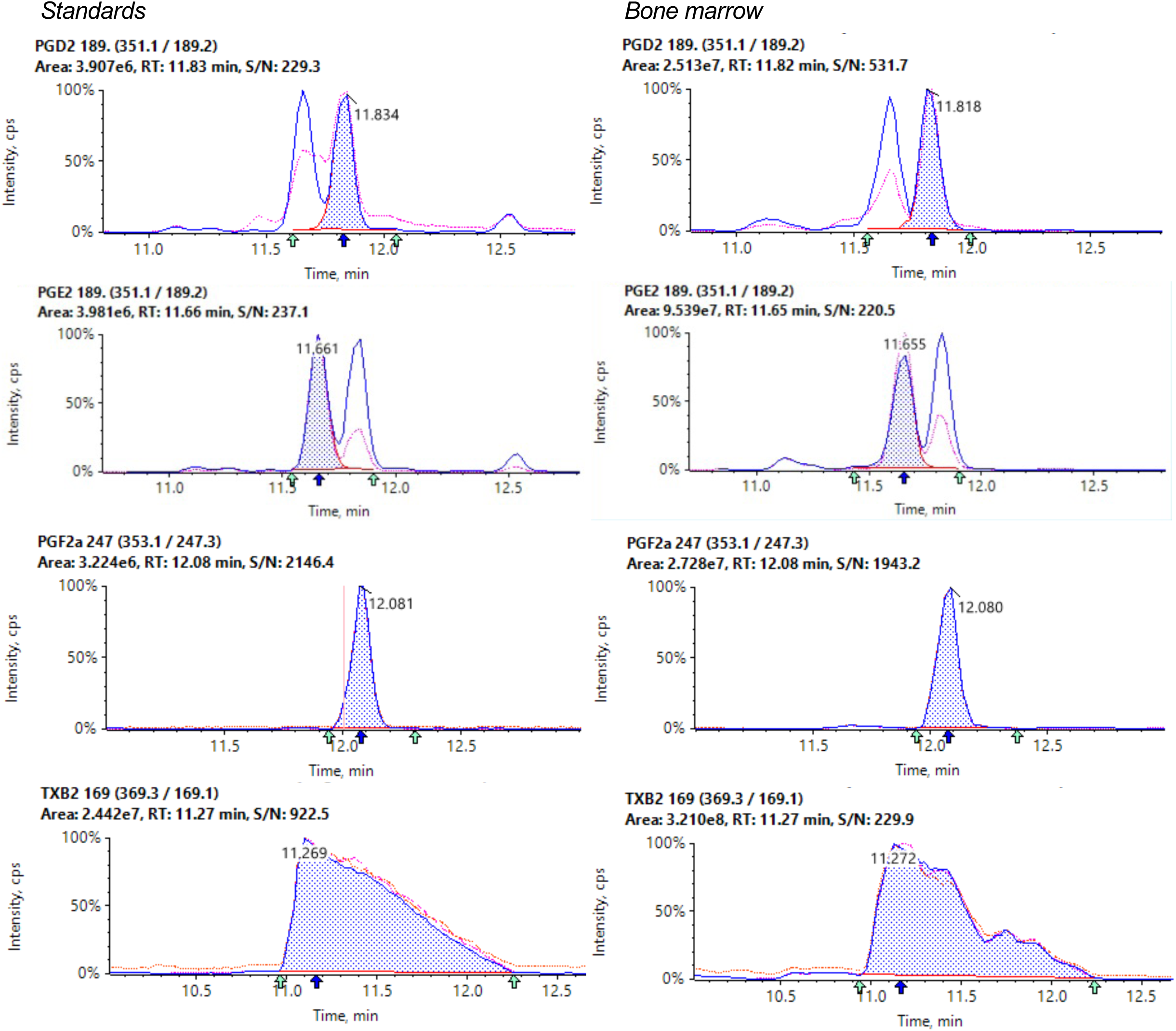
Chromatograms for lipid mediators identified in bone marrows and corresponding standards (ctd) Related to Figure 1A,B

**Figure S2:**
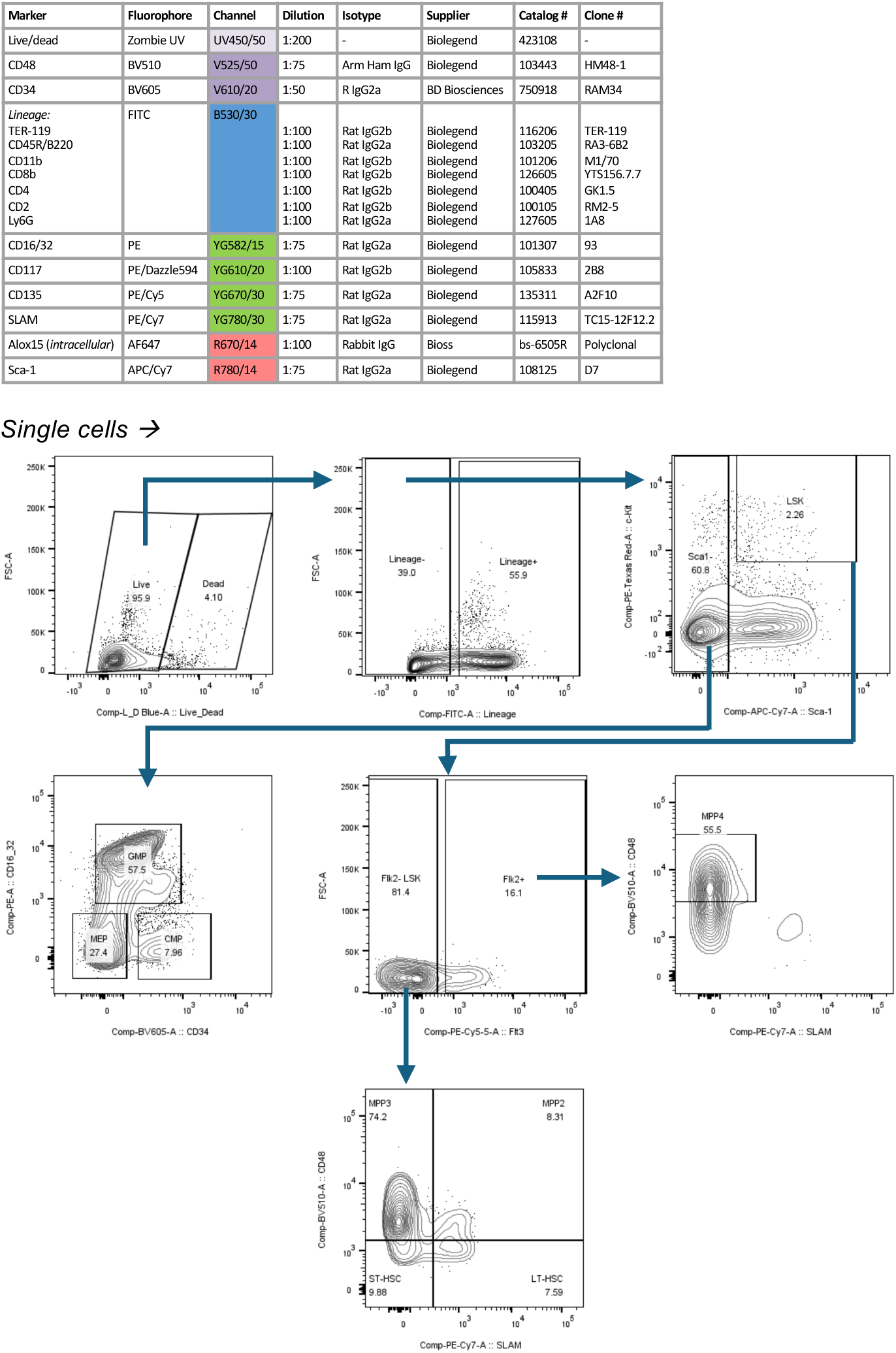
Flow cytometry antibody panels and gating strategies. Related to Figure 1D

**Figure S3:**
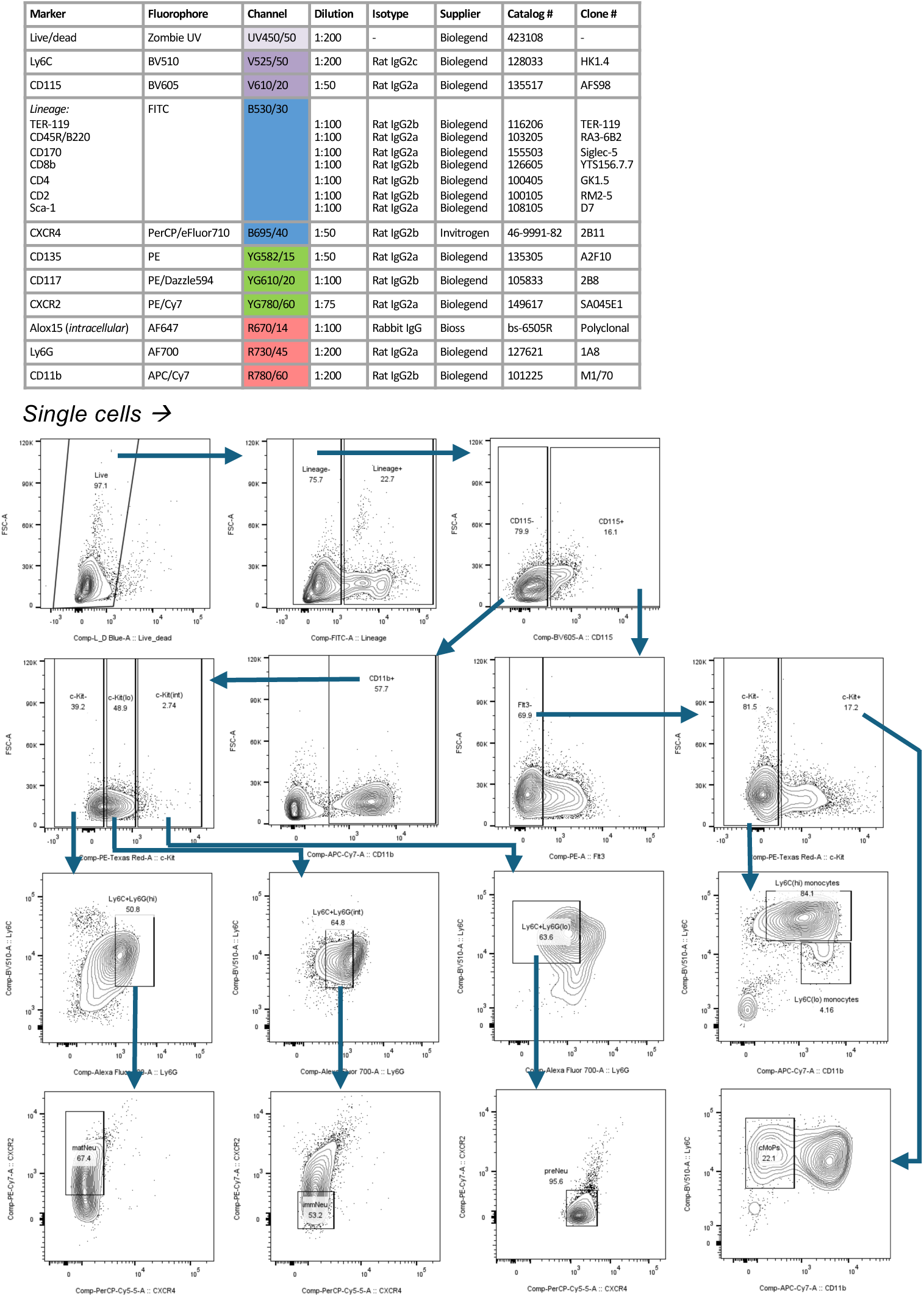
Flow cytometry antibody panels and gating strategies. Related to Figure 1D; Figure 2G-H; Figure 4B

**Figure S4:**
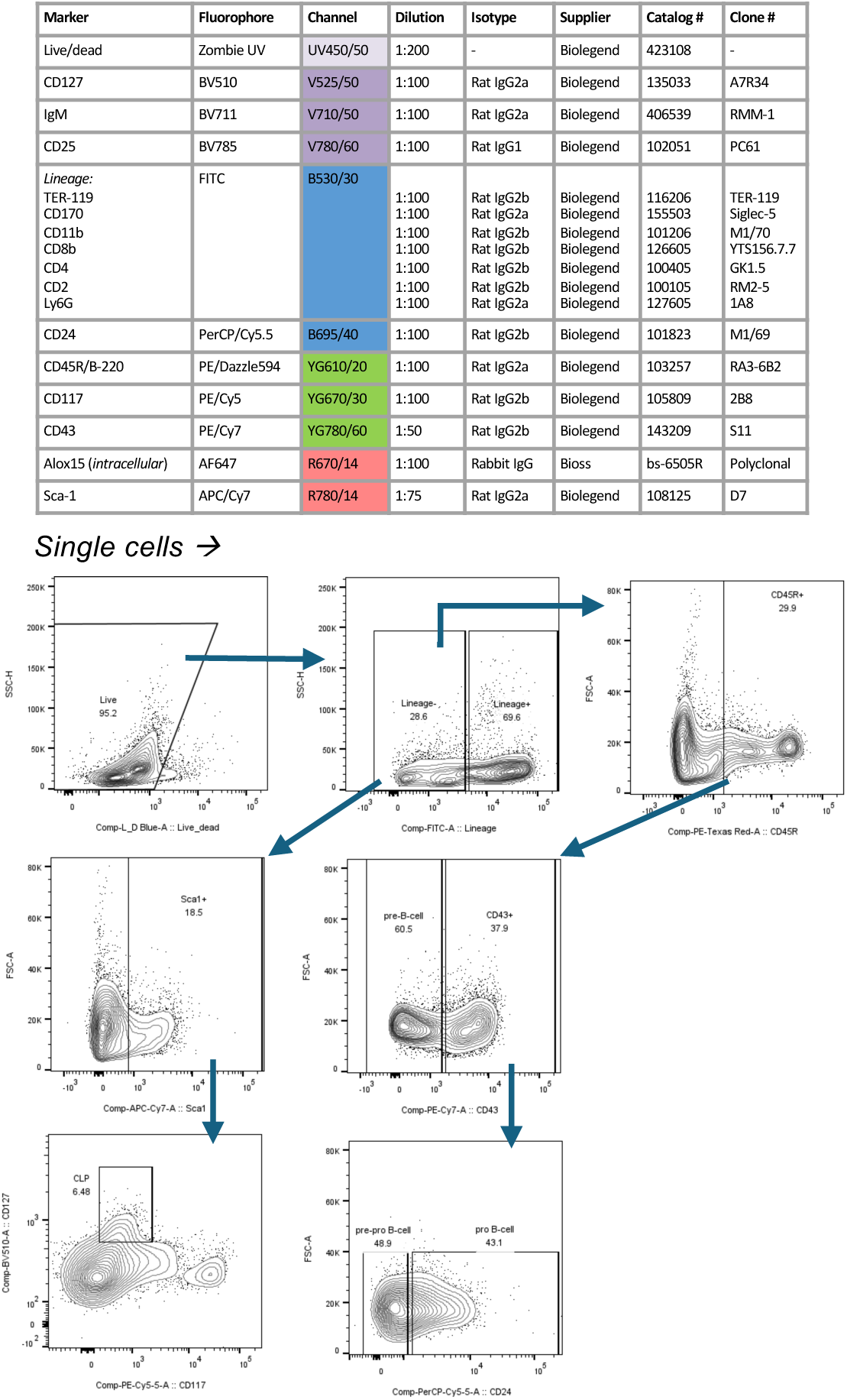
Flow cytometry antibody panels and gating strategies. Related to Figure 1D

**Figure S5:**
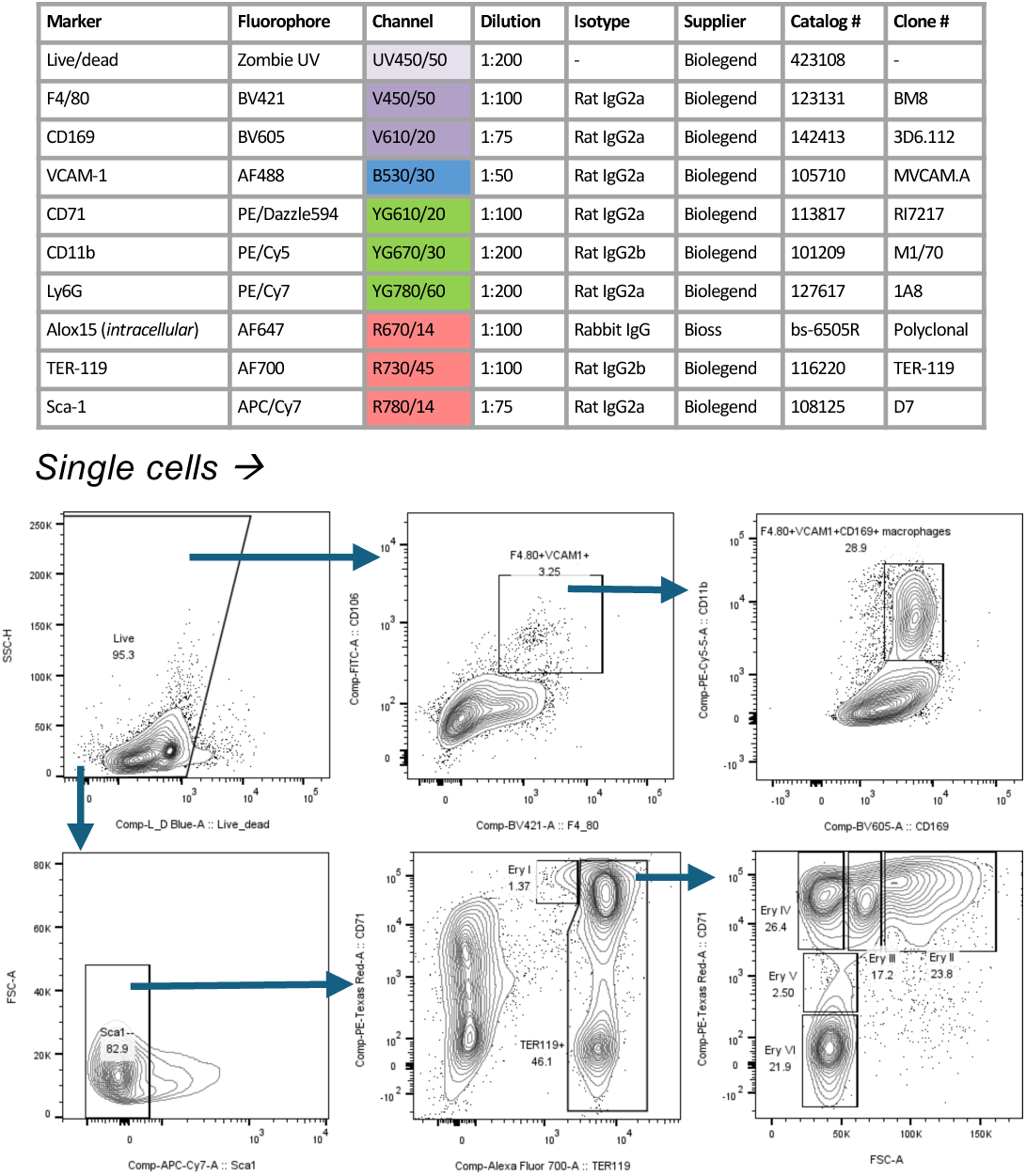
Flow cytometry antibody panels and gating strategies. Related to Figure 1D; Figure 2E-F)

**Figure S6:**
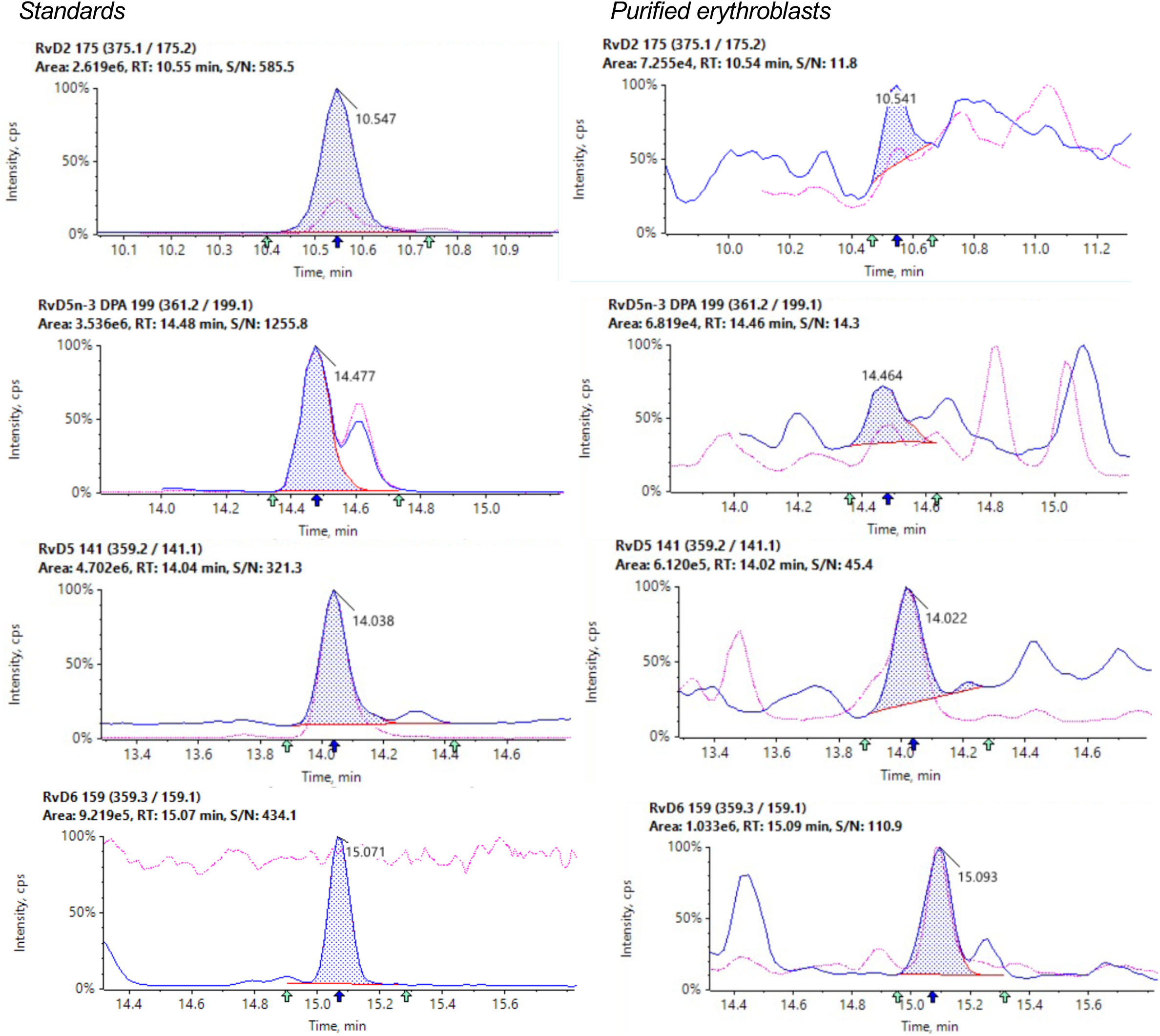
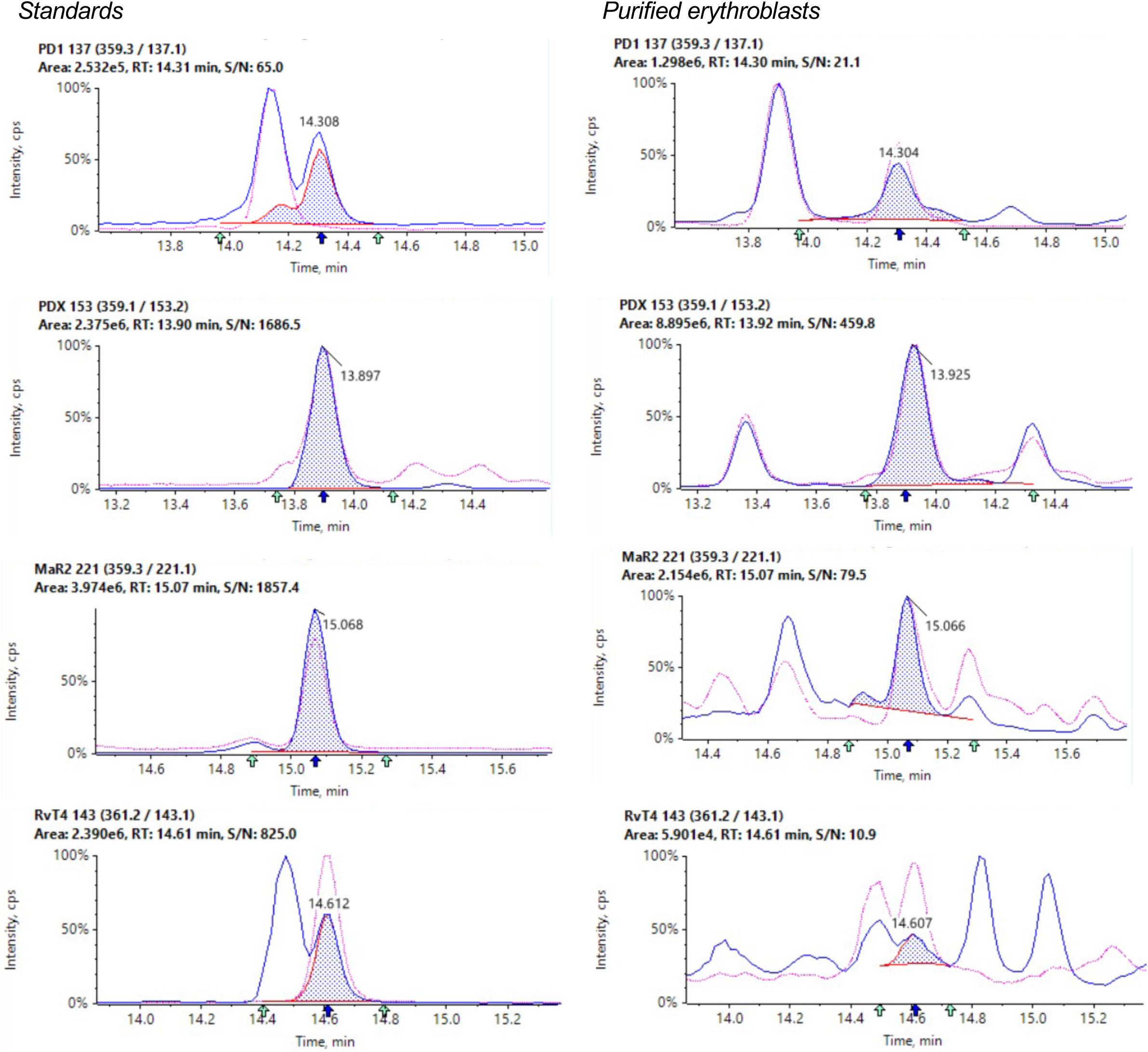
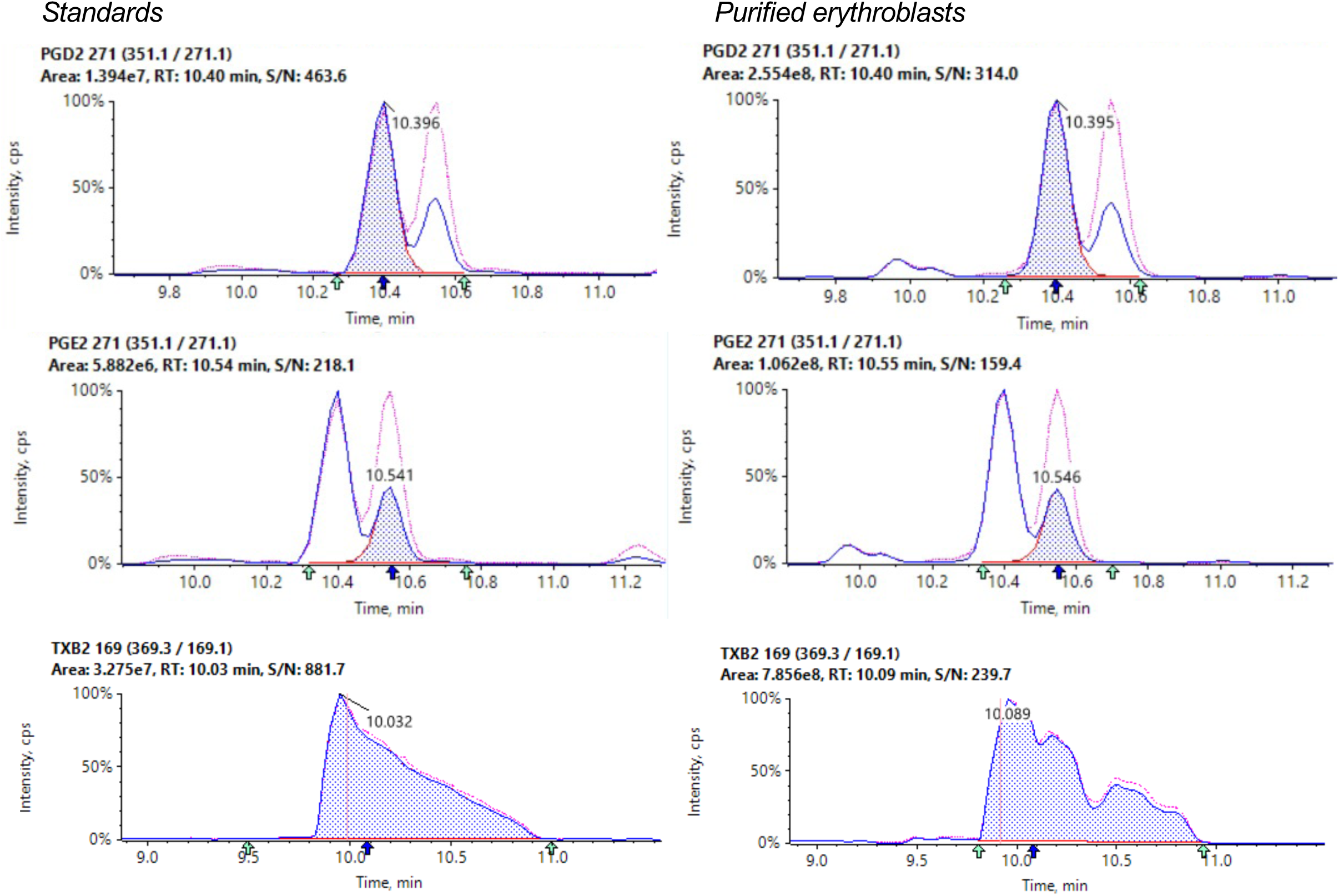
Chromatograms for lipid mediators identified in erythroblasts and corresponding standards. Related to Figure 1G

**Figure S7:**
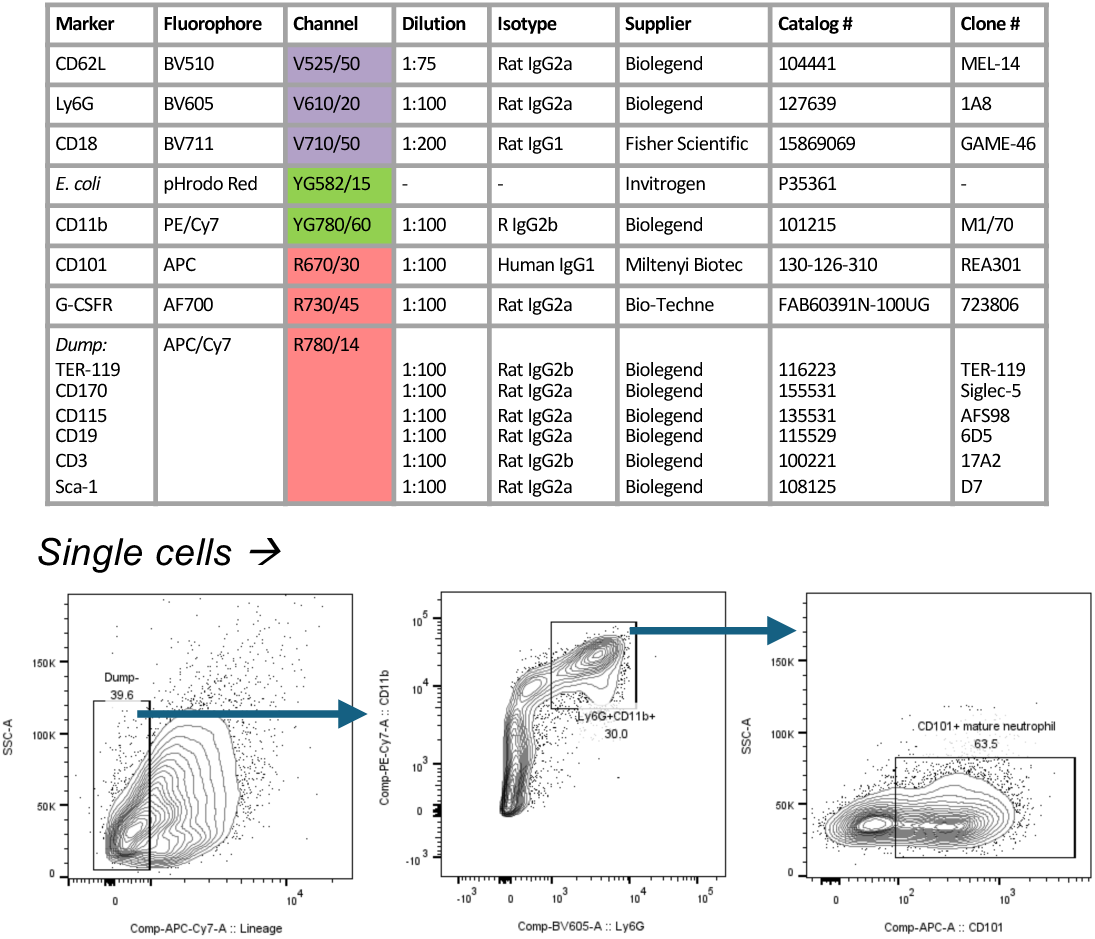
Flow cytometry antibody panels and gating strategies to identify neutrophils in whole blood and ex vivo bone marrow incubations. Related to Figure 3E; Figure 5K; Figure 6H

**Figure S8:**
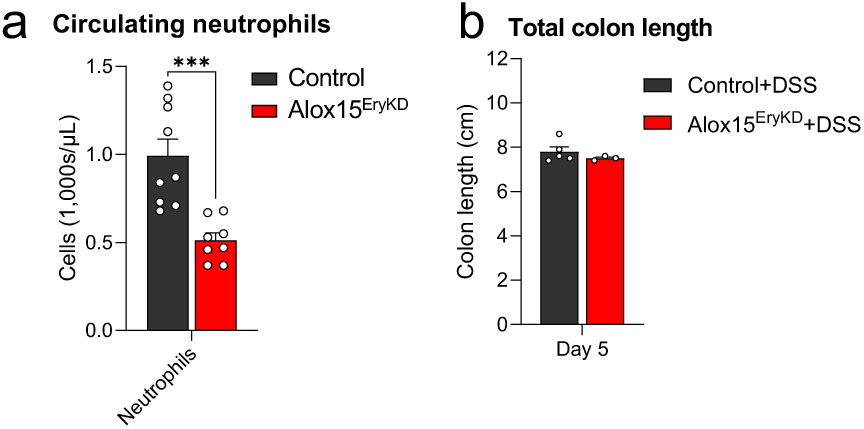
Reduced circulating neutrophil numbers in Alox15^EryKD^ mice. Related to Figure 3 (a) Absolute counts of circulating neutrophils from Alox15^EryKD^ and Alox15-sufficient littermates (Control), n = 9 Control, 8 Alox15^EryKD^ mice. (b) Gross colon lengths at day 5 post-initiation of 2.5% (w/v) Dextran sulphate sodium (DSS)-induced colitis in Alox15^EryKD^ and Alox15-sufficient littermates (Control), n = 5 Control DSS-treated mice, n = 3 Alox15^EryKD^ DSS-treated mice.

**Figure S9:**
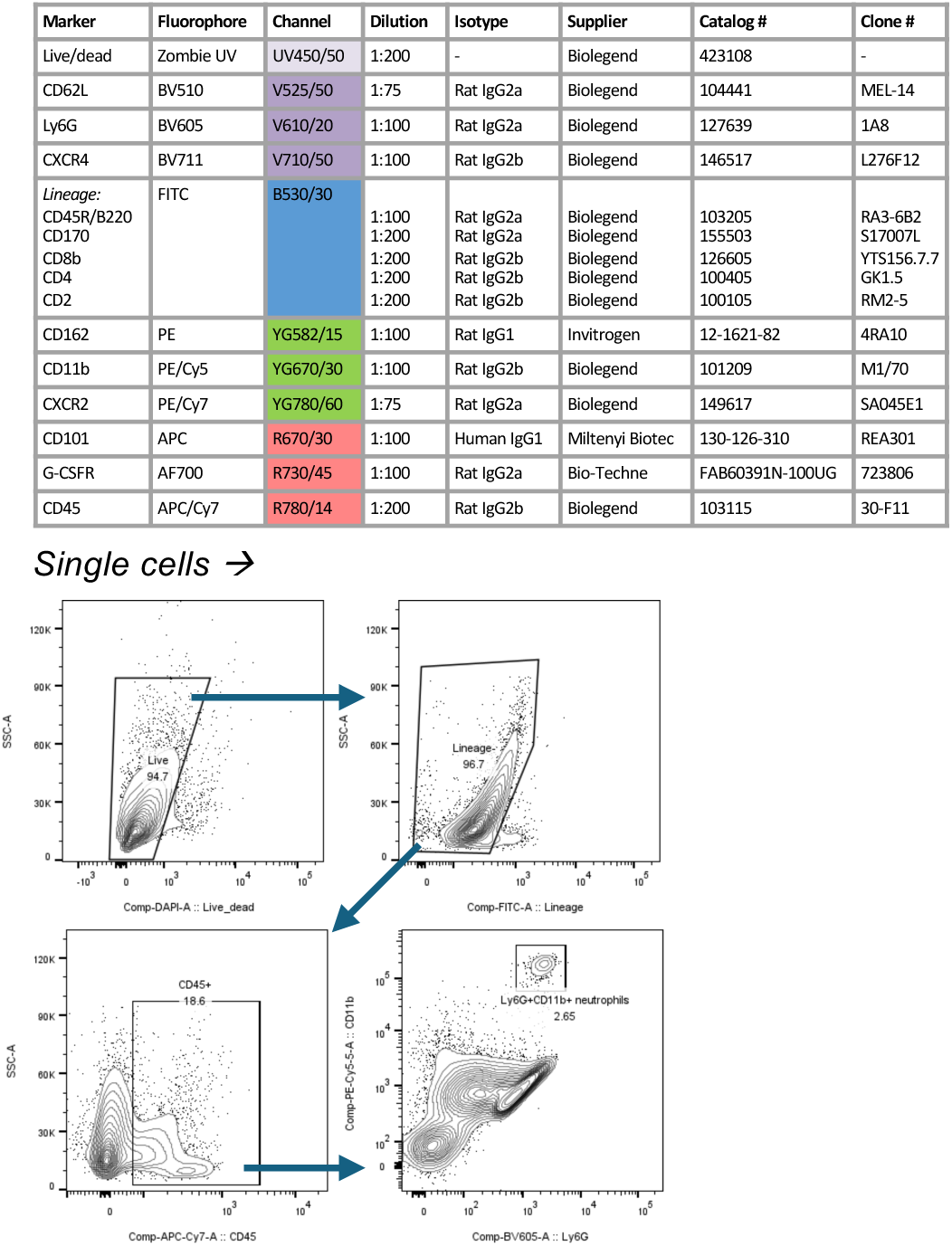
Flow cytometry antibody panels and gating strategies in the identification of neutrophils in peripheral tissues and peritoneal exudates. Related to Figure 3J,N,O; Figure 5I

**Figure S10:**
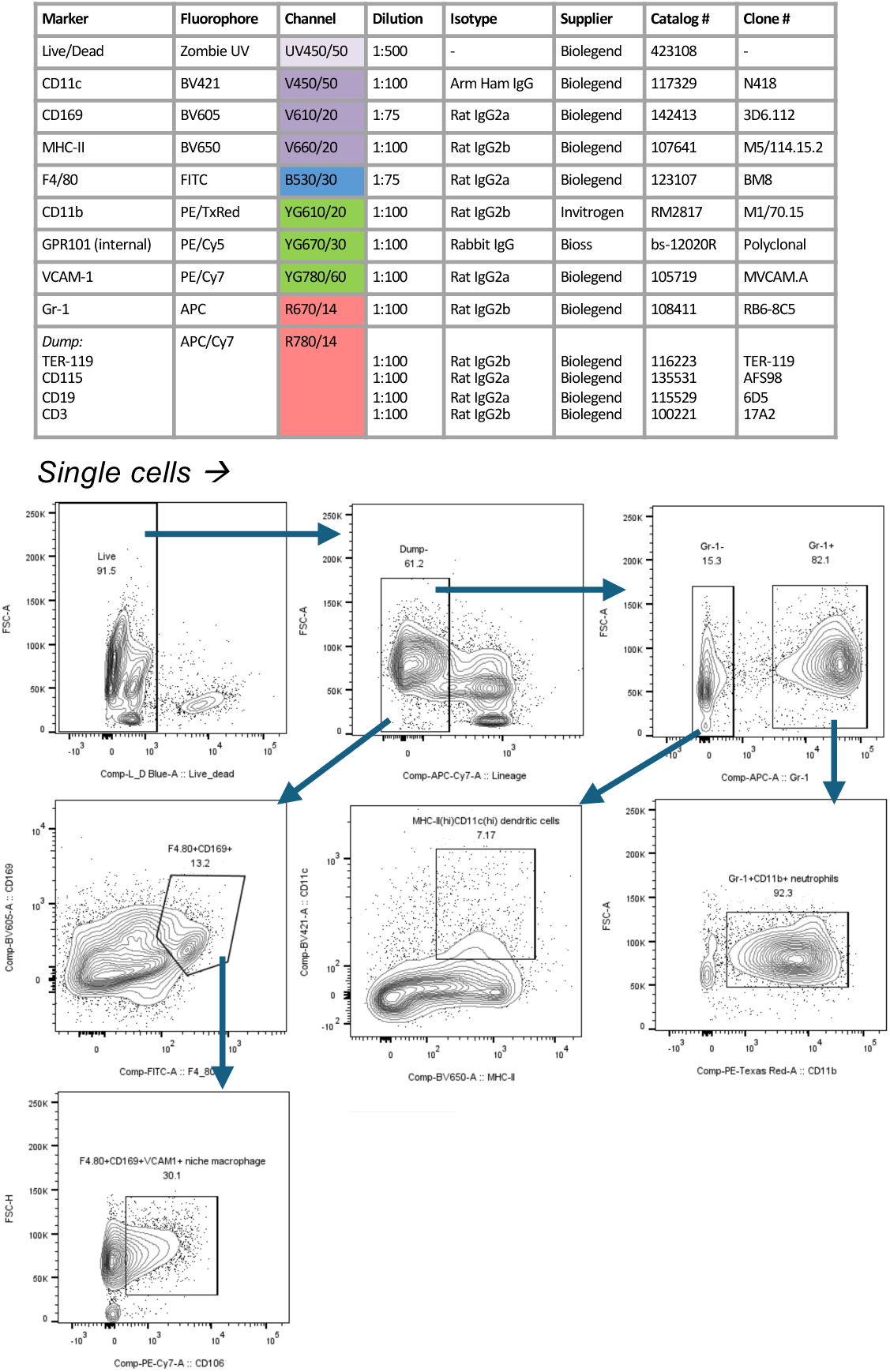
Flow cytometry antibody panels and gating strategies in the identification of macrophages, dendritic cells and neutrophils. Related to Figure 5A, H, I

**Figure S11:**
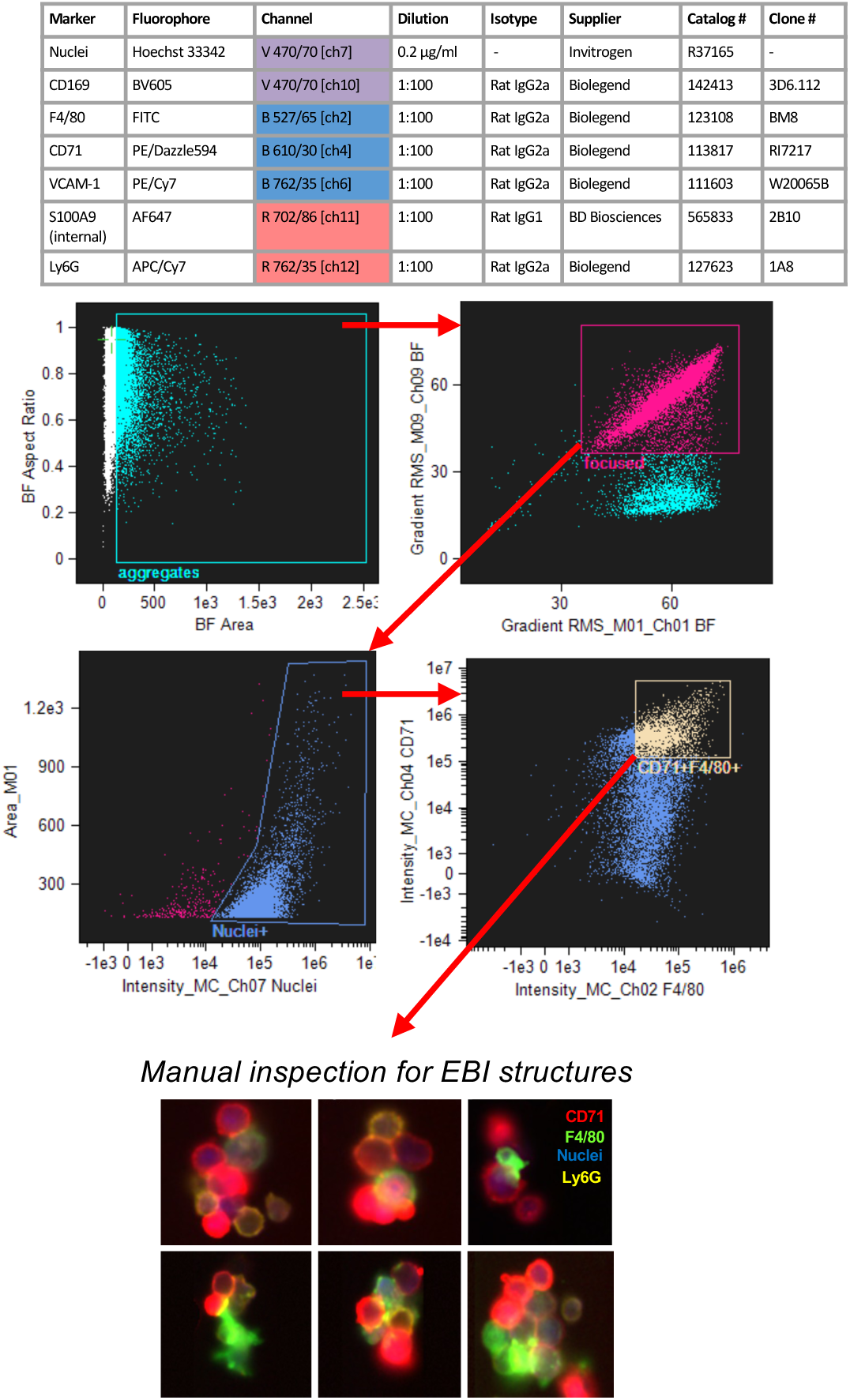
ImageStream antibody panels and gating strategies in the identification of EBIs. Related to Figure 6A-D, F

**Figure S12:**
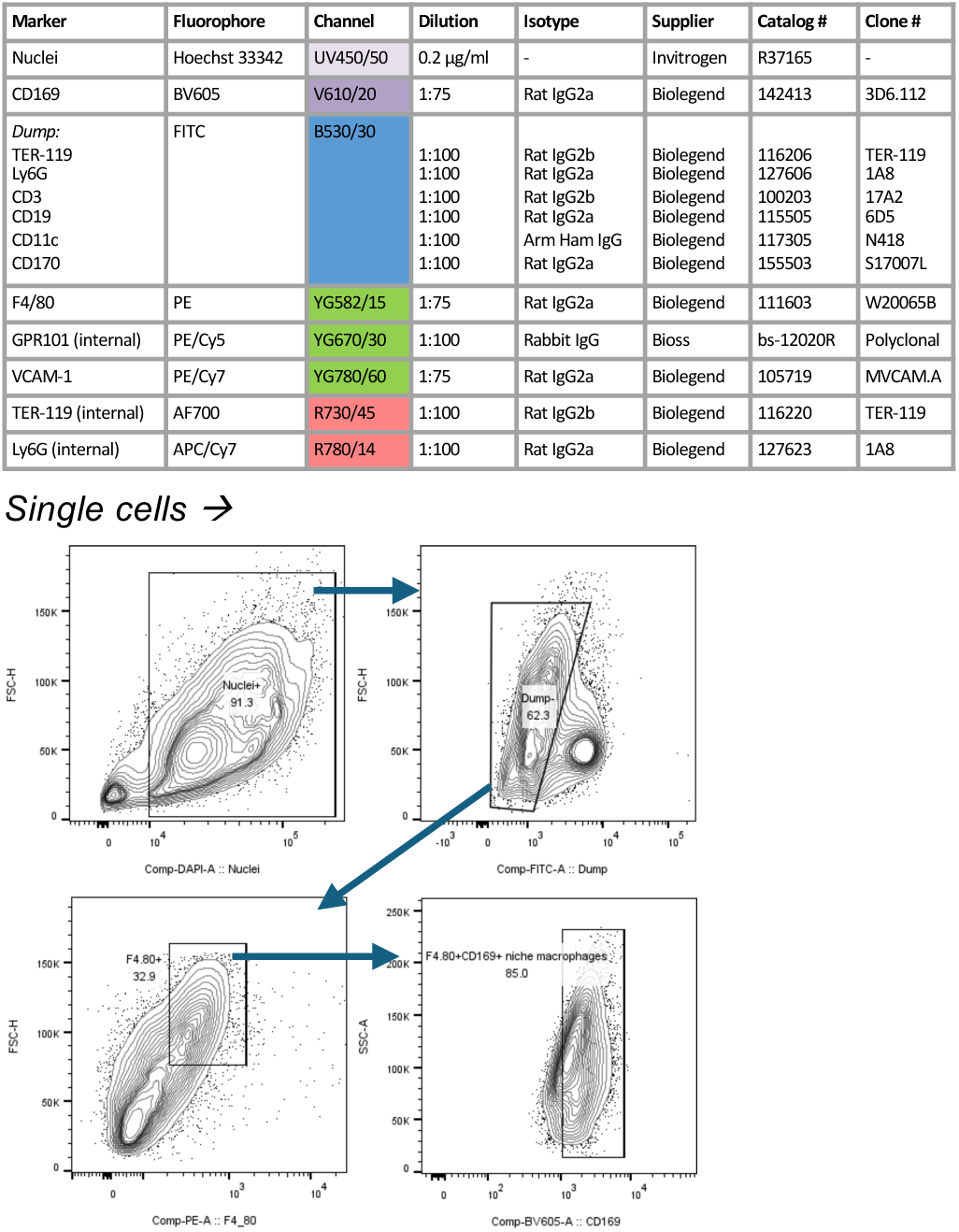
Flow cytometry antibody panels and gating strategies employed in the evaluation of macrophage efferocytosis in ex-vivo cultures. Related to Figure 6G

**Table S1:**
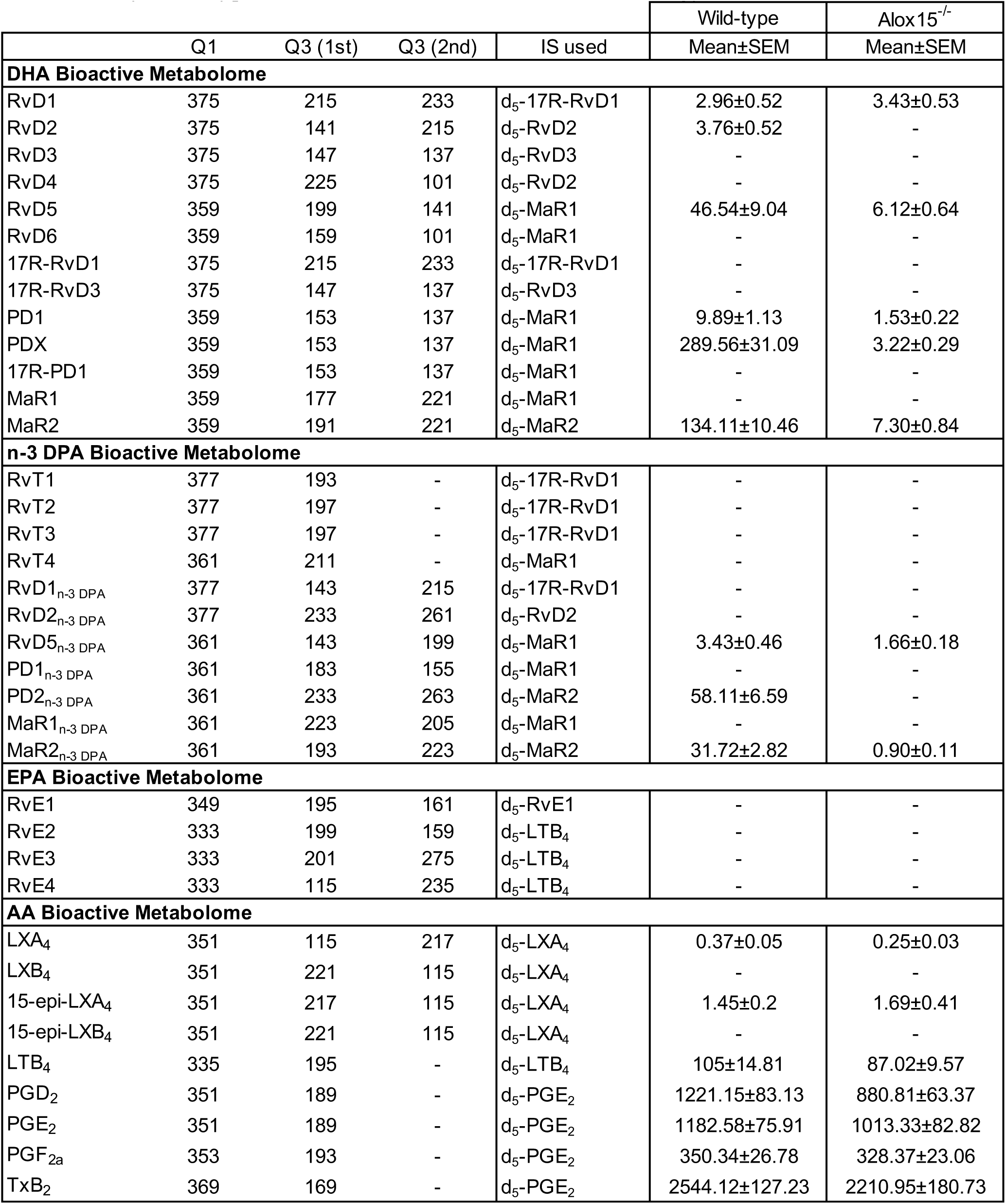
Lipid mediators in bone marrow from Wild-type and Alox15^-/-^ mice - Related to Figure 1A,B Results are reported as pg mediator/20 million bone marrow cells; n = 9 Wild-type, 9 Alox15^-/-^ mice.

**Table S2:**
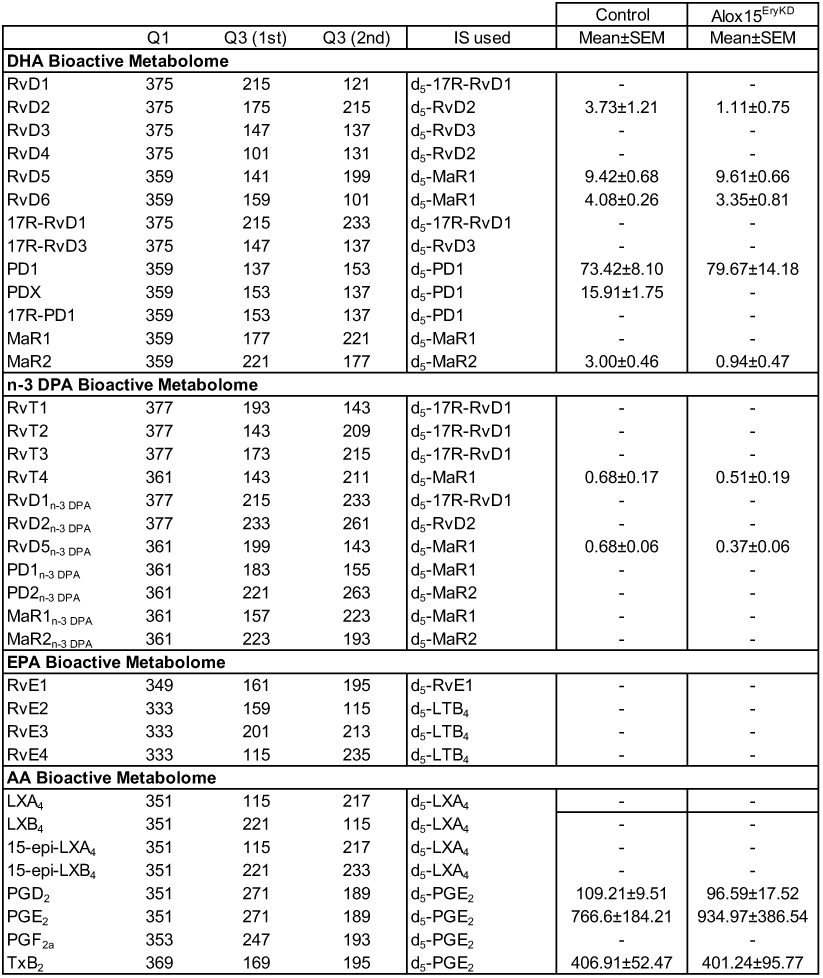
Lipid mediators in purified erythroblasts from Alox15^EryKD^ mice or Control littermates - Related to Figure 1G Results are reported as pg mediator/25 million Ter119+ erythroblasts; n = 6 Control, 7 Alox15^EryKD^ mice

**Table S3:**
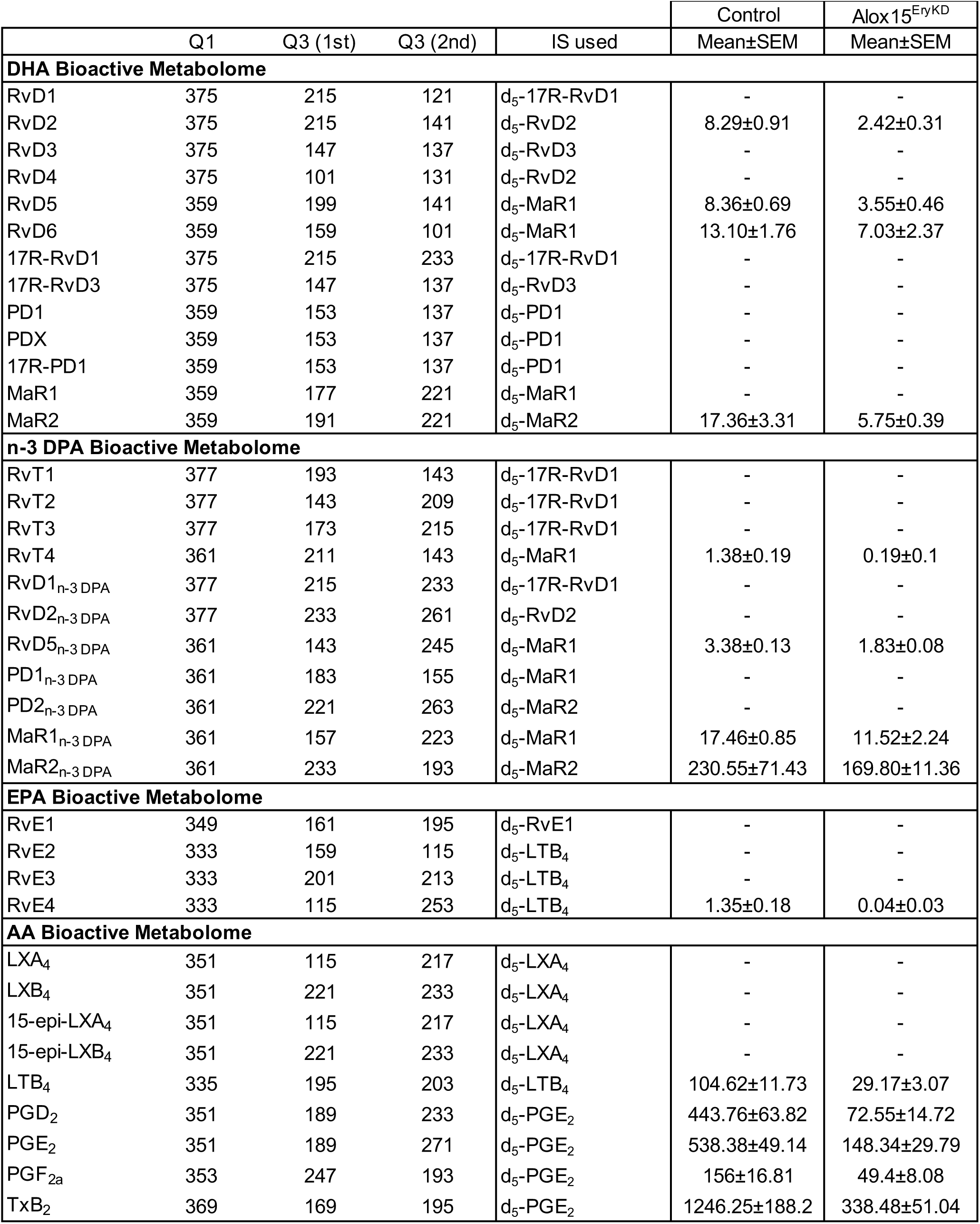
Lipid mediators in bone marrow from Alox15^EryKD^ mice or Control littermates - Related to Figure 1H Results are reported as pg mediator/25 million bone marrow cells; n = 8 Control, 5 Alox15^EryKD^ mice

